# Enhancing inference of differential gene expression in metatranscriptomes from human microbial communities

**DOI:** 10.1101/2025.09.11.675436

**Authors:** Evan M. Lee, Nathan P. McNulty, Matthew C. Hibberd, Jiye Cheng, Kazi Ahsan, Hao-Wei Chang, Barak A. Cohen, Jeffrey I. Gordon

## Abstract

Metatranscriptomic (MTX) sequencing quantifies gene expression from the collective genomes of microbial communities (microbiomes), enabling assessment of functional activity rather than functional potential. While differential expression testing is essential for RNA-sequencing analysis, current metatranscriptomic approaches have only been benchmarked on simulated data, resulting in a lack of standard practices for analysis of real datasets. Here, we use mock communities (defined mixtures of microbial cells with known properties) to quantitatively assess robustness and susceptibility of current approaches to various confounders including organisms’ low relative abundance, differential abundance, low prevalence, global transcriptional output changes, and compositional effects. We show that no current method is robust to all confounders and method performance on simulated data does not generalize to real datasets. We then apply the same approaches to MTX datasets generated from gnotobiotic mice colonized with defined consortia of human bacterial strains and show that the method nominated by the mock community comparisons successfully inferred cross-feeding dynamics that were subsequently validated *in vitro.* Finally, using metagenome-assembled genomes from a human clinical study, we leverage genome-level sequencing depth and detection of genes to exclude low information samples on a per-organism basis to overcome confounding low prevalence and enhance differential expression inference.

## INTRODUCTION

The gastrointestinal tract is home to the largest collection of microbes in our body. Assembly of the gut microbial community (microbiota) begins at birth and progresses over the first several years of postnatal life^1–5^. Culture-independent, DNA-level (metagenomic) analyses have revealed that assembled communities, composed of trillions of microbes from all three domains of life, encode functions not represented in our human genome that are predicted to influence myriad facets of human physiology. Alterations in intestinal community composition have been associated with and causally linked to a variety of disease states^6–20^. The complexity, dynamism and interpersonal variations in community composition manifests over a broad range of temporal and spatial scales^5,21^. These features make the task of defining ‘normal’ formidable^22^. Furthermore, determining the mechanisms that underlie community assembly and robustness as well as host-microbe interactions that contribute to health and disease demands development of methods for analyzing expressed community functions in both non-perturbed and perturbed states.

The decreasing cost of shotgun sequencing of microbial community DNA (metagenomics) has catalyzed an explosion of methods for testing the association of microbes with health and disease states^23–28^, predicting their encoded functions^29–33^, and assembling their individual genomes (metagenome-assembled genomes)^34–37^. These advances enable metatranscriptomic (MTX) sequencing to characterize gene expression of the collective genomes (microbiome) of whole microbial communities, quantifying functional activity more informative than the encoded functional potential inferred from DNA-based analyses^38,39^. The technical challenge of low representation of mRNA in total community RNA has been overcome by commercial kits for depletion of ribosomal RNA (rRNA)^40^, resulting in increasing usage of metatranscriptomic profiling. Efforts comparing method performance to guide tool selection for metagenomic analysis^41–43^ have not yet been extended to metatranscriptomics. Early MTX analyses relied on organism-level RNA:DNA ratios, presence/absence of functions, ordination analyses, taxonomic profiling, and metabolic modeling, which lack statistical hypothesis testing and gene-level resolution^15–20,44^. Consequently, metatranscriptomics has underachieved its potential to characterize gene-level transcriptional regulation of individual organisms within microbial communities.

The identification of genes with statistically significant changes in expression between conditions (differential expression testing) is a fundamental goal of RNA-sequencing (**Fig. 1a**). Single-organism RNA-seq differential expression methods are well-studied and encompass a range of accepted models for identifying gene-level expression changes^45–47^. These methods do not need to account for differences in genome abundance or global transcription rate changes (**Fig. 1b**) because observed sequencing counts are sampled as relative abundances and an organism’s contribution to total RNA is not a factor (**Fig. 1c**). In contrast, any method applied to MTX data must account for the fact that a change in the abundance of one organism within a community affects observed quantification of its genes relative to other species present. Increases in the abundance (DNA-level) and expression (RNA-level) of genes are confounded in metatranscriptomic analyses, which can contribute to false positives if DNA abundance is not controlled for (**Fig. 1d,e**). Metatranscriptomic sequencing data are also heavily zero-inflated for organisms with low prevalence or insufficient coverage due to low relative abundance, resulting in non-detection of genes even without changes in underlying expression (**Fig. 1f,g**).

**Figure 1.**
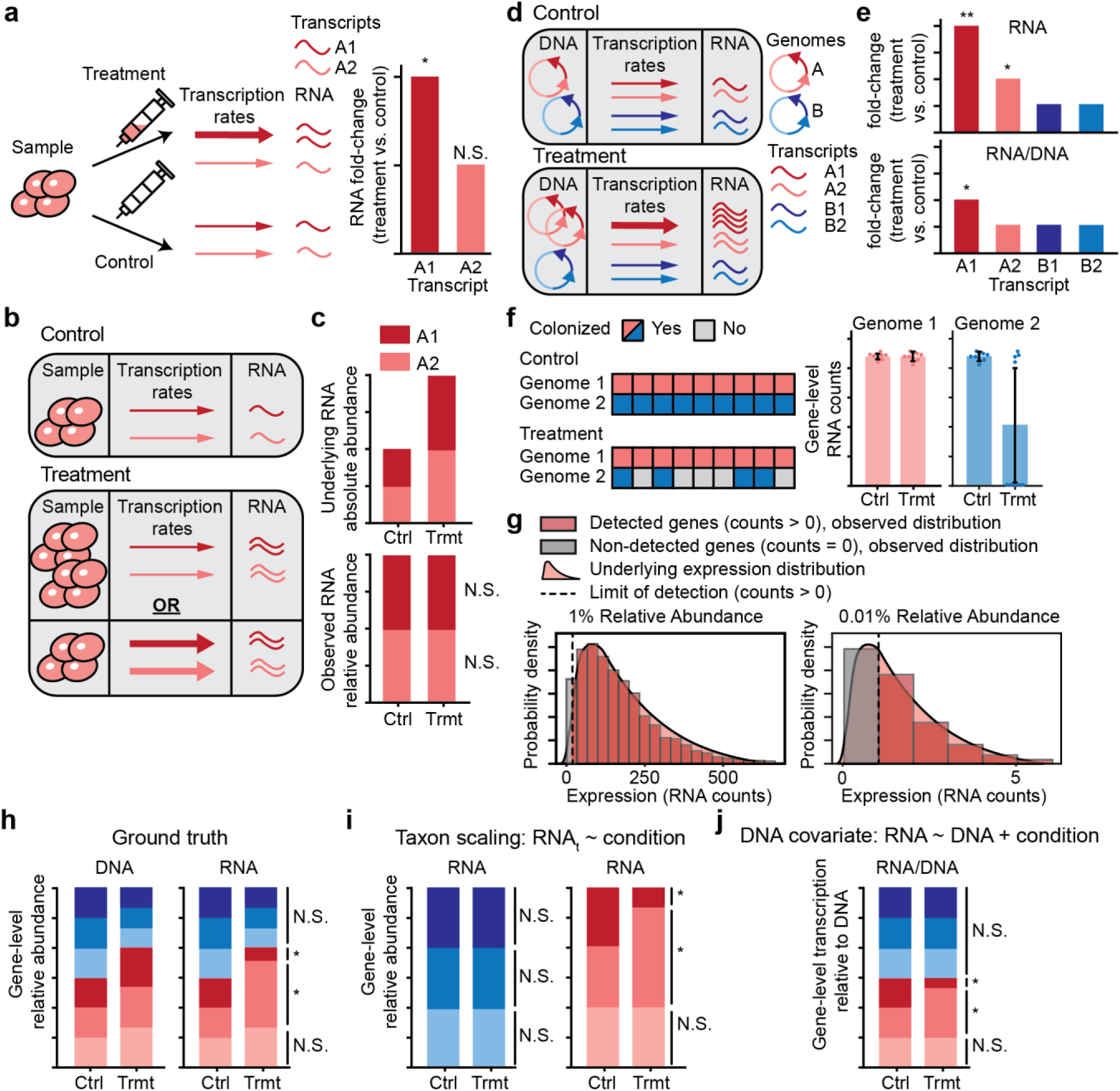
Analytical challenges and existing approaches in metatranscriptomic differential expression. **(a)** Single organism RNA-sequencing and differential expression testing identifies gene-level expression changes between conditions. *: statistically significant differential expression; N.S.: no significant differential expression. **(b,c)** Single organism differential expression is not confounded by increases in genome abundance or global (genome-wide) upregulation (panel b) because although total RNA output is increased, the observed sequencing data are relative abundances and do not quantify underlying RNA absolute abundance (panel c). **(d,e)** In the presence of multiple genomes in a microbial community, increases in genome abundance affect gene-level quantification for metatranscriptomic RNA-sequencing data (panel d), resulting in confounded effects of differential abundance and differential expression for transcript A1 and a false positive result for transcript A2 unless normalizing for DNA abundance (panel e). **(f)** Metatranscriptomic data are zero-inflated due to incomplete prevalence of organism colonization across samples, resulting in zero-inflation and noisy estimation of expression even in the absence of expression changes. **(g)** Insufficient sequencing coverage for organisms at low relative abundance results in an increased fraction of genes below the detection threshold, even in the absence of expression changes. **(h)** Schematic for ground truth DNA and RNA profiles between two conditions, with confounding differential abundance as well as ground-truth differential expression of two genes indicated by asterisks. **(i)** Taxon-specific-scaling controls for changes in genomic abundance by considering genes as relative abundances within each genome individually, allowing selective recovery of differentially expressed genes. RNA_t_: within-taxon RNA abundance. **(j)** Abundance covariate models control for the effects of changes in DNA abundance before considering changes in RNA expression. Abbreviations: Ctrl, control; Trmt, treatment.

Rarefaction (sub-sampling of counts without replacement), historically used for bacterial 16S rRNA amplicon analyses, offers a potential solution to control for changes in genomic representation between conditions. However, the dynamic range of biological variance in organism abundances typically exceeds that of the technical variance in sequencing depths. Rarefaction also inherently compromises statistical power by reducing available information.

The two existing statistical methods for metatranscriptomic differential expression instead rely on directly controlling for genomic abundance dependence during analysis of community expression data. They use either genome-level normalization (‘taxon-specific-scaling’)^49^ or DNA abundance covariates (MTXmodel)^50^ to control for the effects of confounding organism-level differential abundance (**Fig. 1h-j**). A recent effort to comprehensively benchmark available differential expression tools omitted these MTX-specific methods^51^.

Thus far, all reported efforts to assess performance of differential expression methods have relied on simulated data^49–51^, where assumptions made during synthetic data generation can influence results and artificially inflate method performance^52^. Consequently, differential expression analysis of metatranscriptomes lacks standard practices and quantitative assessment of the performance and biases of existing approaches^53^. Nonetheless, existing tools have been applied broadly and enabled discovery of genes involved in polyphenol metabolism^54^ and adaptions to mucosal inflammation by opportunistic pathogens^55,56^. These efforts highlight both the potential of and continuing need for differential expression tools suitable for microbiome applications.

Mock microbial communities are critical for assessing the performance of analytical approaches in microbiome studies^52^. These communities contain mixtures of microbial cells in known ratios and thus enable systematic assessment of method performance (benchmarking) in recovering the ground truth composition. While they are limited by low sample diversity and the labor and cost involved in creating them, they represent a gold standard for performance comparisons under real operating conditions, including technical sequencing and biological noise^52^. They have been employed to benchmark 16S rRNA amplicon, shotgun metagenomic, and cytometric analyses of community composition^55–61^, but there are no reports of such datasets being used to evaluate differential expression tools for microbial communities, partly due to the difficulty of knowing biological ground truth regarding gene regulation.

In this study, we first perform method benchmarking on simulated data and demonstrate the low sensitivity of existing methods for identifying differentially expressed genes from organisms present at low relative abundance. We then employ mock communities to evaluate differential expression methods on real bacterial datasets by benchmarking against a gold-standard single-organism transcriptomics method (DESeq2^45^) applied to samples from cultures containing a single organism. Using simple comparisons with emulated confounding features such as organism-level low relative abundance, differential abundance, low prevalence, global transcription rate changes, and compositional effects, we demonstrate the robustness and susceptibility of different methods to the inherent challenges of MTX datasets, show that method performance on simulated data does not generalize to real samples, and provide recommendations for sequencing effort and analytical approaches for adequate differential expression testing. We then apply these methods to published datasets generated from (i) gnotobiotic mice colonized with defined consortia of human gut bacterial strains and (ii) a clinical study involving serial sampling of the fecal microbiota of participants before, during and at the end of administration of a microbiota-directed therapeutic food. We conclude that MTX method benchmarking on real, not simulated, datasets can and should guide methodological choice and implementation, enabling inference and validation of microbial metabolic strategies and interactions from *in vivo* datasets.

## RESULTS

### Benchmarking on simulated data

As noted above, simulated data are advantageous for method benchmarking due to their inexpensive and straightforward production, high community complexity, flexible properties, and explicit definition of ground truth^52^. Previous studies have not compared all reported MTX-specific tools, so we first adapted the benchmarking strategy used for MTXmodel^50^ and extended it to additional methods^49,51^. Six published simulated datasets, each with 100 samples split into cases and controls, are described in **Fig. 2a** and **Supplementary Table 1a**; they include a ‘null’ dataset without simulated differential expression but with confounders (e.g. differential abundance), as well as ‘true’ datasets with simulated differential expression of varying strengths or in combination with confounders. For each dataset, DNA and RNA counts were randomly sampled from log-normal distributions parameterized using empirical prevalence and abundance data from the Human Microbiome Project^62^.

**Figure 2.**
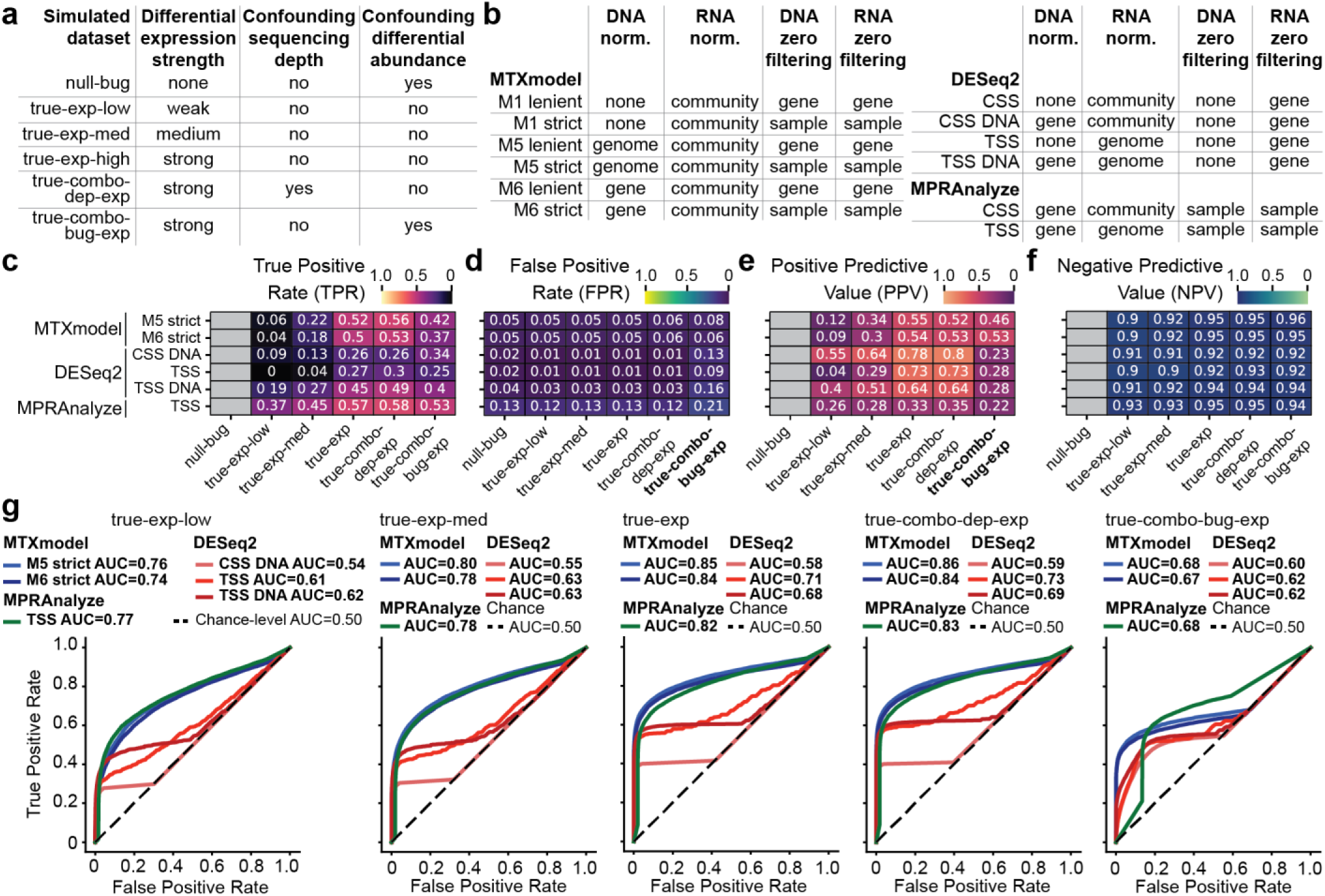
Statistical differential expression method benchmarking on simulated datasets. **(a,b)** Characteristics of simulated datasets (panel a) and differential expression methods and implementations tested (panel b). **(c-f)** True positive rate (TPR, or sensitivity; panel c), false positive rate (FPR, or 1-specificity; panel d), positive predictive value (PPV, or precision; panel e), and negative predictive value (NPV; panel f) for MTXmodel, DESeq2, and MPRAnalyze across a range of parameter choices in six simulated datasets. Implementations and datasets specifically emphasized in the text are bolded. Metrics that are not defined for datasets lacking ground truth differential expression are grayed out. (**g**) Area under the curve (AUC) quantification and receiver operating characteristic curves for statistical methods in the five synthetic ‘true-’ datasets which included simulated differential expression. An AUC of 0.5 indicates the classification power of random guessing, with greater AUC indicating a higher ability to discriminate differentially expressed and non-differentially expressed genes. Abbreviations: M1, no DNA abundance covariates; M5, genome-level DNA covariates; M6, gene-level DNA covariates; CSS, community-sum-scaling; TSS, taxon-specific-scaling; DNA, DNA abundance normalization.

Calculation of true positive rate (sensitivity) and false positive rate (1 - specificity) were performed as described previously^50^, and additional calculations of positive predictive value (precision) and negative predictive value were applied (see *Methods*). These metrics respectively quantify successful recovery of differentially expressed and non-differentially expressed genes, and the reliability of significant and non-significant differential expression results. An ideal method would maximize sensitivity, precision, and negative predictive value, as well as minimize false positive rate, thereby maximizing specificity.

Each differential expression method was initially run under a variety of parameters to determine those that optimized its performance (**Fig. 2b**, **Supplementary Fig. 1**; see *Methods*); the methods which performed best by our metrics are shown in **Fig. 2c-g**, while all methods tested are shown in **Supplementary Fig. 2**. This analysis included the following methods and implementations: (i) DESeq2^45^, a gold-standard negative binomial model for single-organism RNA-seq. We used DESeq2 with either default community-sum-scaling (CSS, equivalent to community-level total-sum-scaling^50^) or modified with taxon-specific scaling^49^ (TSS), where sequencing depth normalization occurs for all genomes together or for each genome individually. We additionally implemented abundance covariates for DESeq2 by including DNA abundances as gene-level normalization factors (‘CSS DNA’ or ‘TSS DNA’); (ii) MTXmodel^50^, a widely used MTX-specific approach that incorporates DNA abundances as covariates in log-normal models. We used either no abundance information (model M1 in Zhang et al.^50^), or genome- or gene-level abundances (models M5 and M6, respectively). We also tested their ‘strict’ and ‘lenient’ filtering of zeros, which respectively exclude samples or genes that have zero DNA or RNA counts (see *Methods*) - of these, the authors recommend gene-level covariates and ‘strict’ filtering; (iii) MPRAnalyze^63^, a gamma and negative binomial model for DNA and RNA, respectively. MPRAnalyze was designed for massively parallel reporter assays which quantify expression relative to abundance, similar to the analytical challenge of metatranscriptomics. We utilized MPRAnalyze with either community- or taxon-level scaling.

Consistent with reported results^50^, MTXmodel exhibited highest sensitivity when using genome- or gene-level DNA abundance and ‘strict’ sample-level filtering of zeros (respectively, ‘M5 strict’ and ‘M6 strict’) (**Fig. 2c**). Of all MTXmodel implementations, gene-level DNA abundances exhibited the best control of false positives in the presence of confounding differential abundance (‘true-combo-bug-exp’; **Fig. 2d**). Taxon-scaling improved both sensitivity and specificity for DESeq2 and increased sensitivity but lowered specificity for MPRAnalyze (**Supplementary Fig. 2c,d**). Because of improved false positive control, DESeq2 with either DNA covariates (CSS DNA) or taxon scaling (TSS) exhibited the highest precision of all methods tested except in the presence of differential abundance, indicating the highest reliability of its reported significantly differentially expressed genes (**Fig. 2e**). All methods had similar negative predictive value due to the high representation of non-differentially expressed genes (as high as 90% of the datasets) (**Fig. 2f**).

To assess each model’s ability to balance sensitivity and specificity across varying significance thresholds, we turned to receiver operating characteristic (ROC) area under the curve (AUC) quantification (see *Methods*). Across the five datasets with simulated differential expression, MTXmodel and taxon-scaled MPRAnalyze, the two methods leveraging DNA abundances, exhibited the highest AUC (**Fig. 2g, Supplementary Fig. 2g**). Because there were not substantial differences in performance for MTXmodel using genome- or gene-level DNA abundance (M5 strict and M6 strict, respectively), we adhered to gene-level covariates as recommended by Zhang et al. for remaining analyses. Notably, despite the high precision of their predictions, DESeq2 CSS DNA and TSS DNA exhibited lower AUCs than taxon-scaled DESeq2 without DNA covariates due to having a larger fraction of missing differential expression predictions (**Fig. 2g**; see *Methods*). Based on these results, we advanced MTXmodel implementations ‘M6 strict’ and ‘M6 lenient’, DESeq2 implementations CSS DNA and TSS, and MPRAnalyze implementation TSS as promising candidates for further analyses. The community-scaled versions of the latter two methods (DESeq2 CSS, MPRAnalyze CSS) were also advanced as controls.

We hypothesized that low sequencing coverage of organisms with low relative abundance would lead to decreased sensitivity and precision for identifying their differentially expressed genes. To test this, the 100 organisms in these simulated datasets were binned into deciles by their ranked relative abundance (see *Methods*). We then calculated sensitivity, specificity, and precision for each decile separately (**Supplementary Fig. 2h-j**). All methods exhibited low sensitivity and precision for genes from organisms in the bottom four deciles, corresponding to a relative abundance of approximately 0.01% or below (**Supplementary Fig. 2h,j; Supplementary Table 1c,d**).

In these datasets, MTXmodel outperformed other methods across a variety of metrics and especially in the presence of confounding differential abundance. However, the assumptions of log-normal distributed counts and linear scaling of RNA expression with DNA abundance made during synthetic data generation align with the statistical models used by MTXmodel, warranting further assessment of the generalizability of these results. Additionally, these results indicate that current reported methods cannot overcome the limitation of decreased information due to low sequencing coverage for organisms at low relative abundance. This is consistent with prior evidence describing insufficient resolution for single transcript analyses for low abundance species^64^. To test the generalizability of method performance and the effects of low abundance in real datasets, we turned to mock community samples which provide ground-truth composition, allow inference of differential expression through gold-standard methods, and incorporate biological and technical noise seen under real operating conditions.

### Benchmarking using *in vitro* mock communities

Simulated counts are a powerful tool for assessing method performance because they allow precise control over dataset properties (e.g., strength of differential expression) and critically provide ‘ground truth’ of which genes are differentially expressed. However, they are limited in that simulation model choices implicitly make assumptions about data generation that may bias certain methods’ performance and may not generalize to real sequencing datasets^52^. In order to determine which methods achieved the highest sensitivity and specificity on real microbial expression data and whether their performance generalized from tests using simulated counts, we prepared mock communities with well-defined properties. These two-member mock communities were assembled using *Prevotella (Segatella) copri* grown in a defined medium with either 1% arabinan or 1% glucose as the sole carbon source. *P. copri* from either growth condition was then mixed in predefined ratios with a ‘background community’ composed of *Escherichia coli* to emulate specific relative abundances (n=6 per growth condition per relative abundance; **Fig. 3a; Supplementary Table 2**). We reasoned that despite the low species richness of these mock communities, method performance would generalize to more complex communities so long as sequencing reads are accurately assigned to each organism.

**Figure 3.**
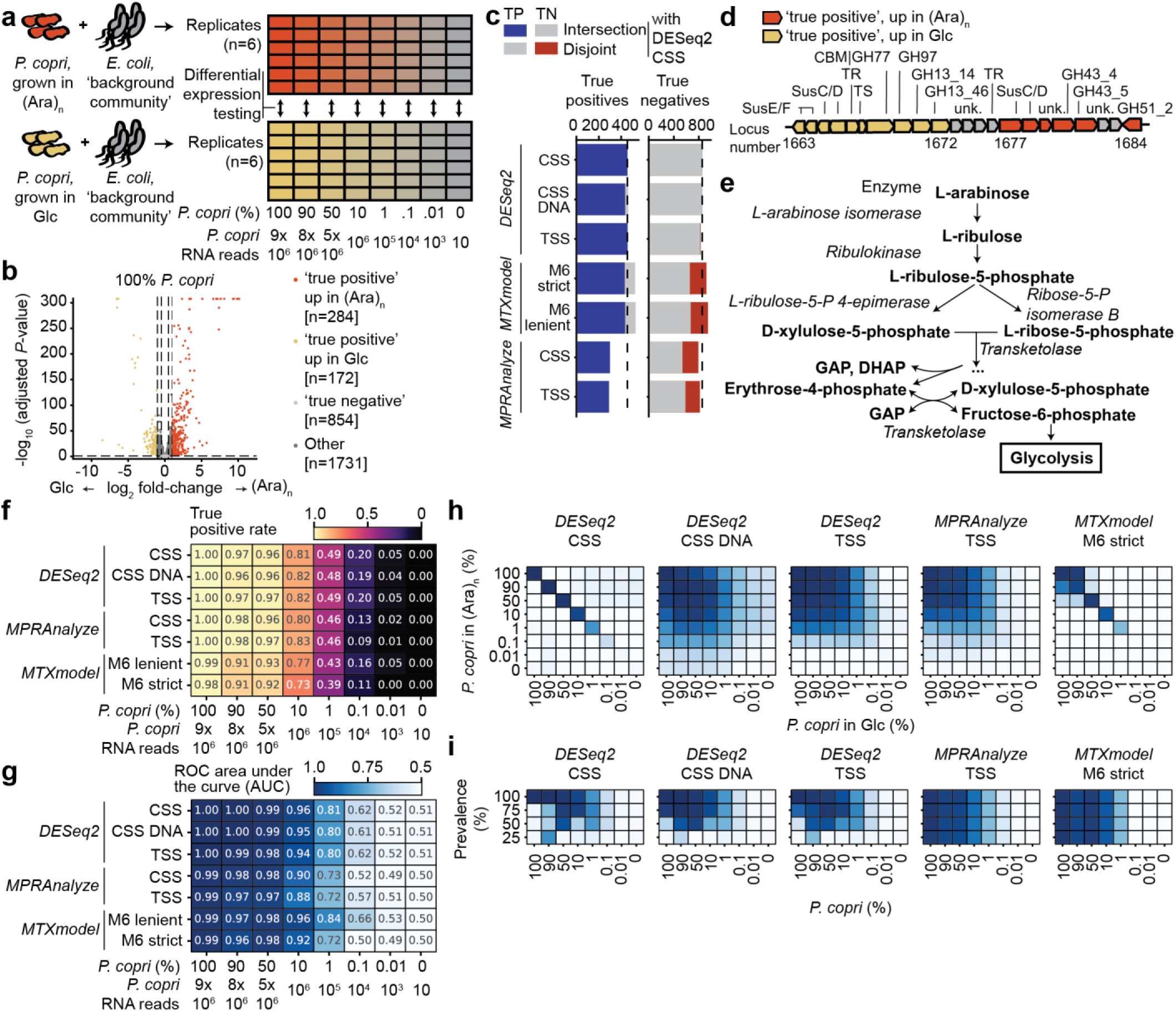
Statistical method benchmarking on datasets from mock communities. **(a)** Experimental design. **(b)** ‘True positive’ and ‘true negative’ gene sets were defined by applying a gold-standard method, community-scaled DESeq2, to the single-organism 100% *P. copri* samples in arabinan and glucose. Vertical dotted lines indicate the log_2_ fold-change thresholds for ‘true positive’ and ‘true negative’ at ± 1 and ± 0.5 respectively, while the horizontal line shows the adjusted *P*-value threshold of 0.05. Sizes of the ‘true positive’ and ‘true negative’ gene sets, along with the number of genes with ambiguous differential expression evidence that are not part of either of these sets are shown in the legend. (**c**) Number of genes in the intersection (overlapping with) and disjoint (unique to) sets when comparing ‘true positive’ and ‘true negative’ sets defined by other benchmarked methods compared to those defined by community-scaled DESeq2. Dashed lines indicate the number of genes defined by community-scaled DESeq2. (**d**) A predicted polysaccharide utilization locus (PUL) with annotated carbohydrate active enzymes, transcriptional regulators, and SusC/D glycan-binding and transporter pairs in the *P. copri* genome containing the most strongly differentially expressed genes that were upregulated by glucose (yellow) or arabinan (orange). (**e**) Schematic of *Prevotella* pentose-phosphate pathway catabolism of arabinose with enzymes that were significantly upregulated in arabinan italicized. (**f, g**) True positive rate (TPR; panel f) and receiver operating characteristic (ROC) area under the curve quantification (AUC; panel g) for benchmarked methods when analyzing the different mixture ratios of *P. copri*, from highest to lowest relative abundance. (**h**) ROC AUC for methods in comparisons with varying strengths of differential abundance, emulated by performing all pairwise comparisons between mixture ratios. (**i**) ROC AUC when emulating decreasing prevalence by including either 0, 2, 6, or 18 100% *E. coli* samples with the six *P. copri*-containing samples in each group. Abbreviations: (Ara)_n_, arabinan; Glc, glucose; TP, true positive; TN, true negative; CSS: community-sum-scaling; CSS DNA: community-sum scaling with DNA abundance normalization; TSS, taxon-specific-scaling; TPR, true positive rate; FPR, false positive rate; PPV, positive predictive value; GH, glycoside hydrolase; CBM, carbohydrate-binding module; SusC/D, TonB-dependent transporter (SusC) and surface glycan-binding protein (SusD); TR, transcriptional regulator; TS, other transporter; unk, unknown function; GAP, glyceraldehyde 3-phosphate; DHAP, dihydroxyacetone phosphate; ROC, receiver operating characteristic; AUC, area under the curve.

*P. copri* is a saccharolytic bacterium that possesses diverse carbohydrate utilization machinery and is prevalent in non-Westernized populations^63,64^. Its ability to degrade complex plant glycans and to produce succinate has been associated with beneficial host effects in the contexts of insulin resistance and childhood undernutrition^65–69^, although its metabolic activities downstream of carbohydrate utilization remain poorly characterized. *E. coli* was selected to represent a background community based on its taxonomic distance from *P. copri*, allowing accurate mapping of sequencing reads to each organism. Both strains in the mock communities had been isolated previously and complete, high-quality long-read genome assemblies generated^70^, allowing reliable and specific gene-level quantification. We reasoned that DESeq2, a gold standard-differential expression method for single-organism transcriptomics, could be used to identify the ‘ground truth’ differentially expressed genes inferred from samples containing only *P. copri* exposed to either carbon source (**Fig. 3b**). Because the two organisms were grown separately and transcriptionally ‘frozen’ prior to being mixed, the same set of differentially expressed *P. copri* genes should be identifiable in both the monoculture and mixture contexts. The performance of other methods could then be benchmarked against this initial comparison while introducing confounding factors like low relative abundance, differential abundance, and non-colonization.

We characterized and confirmed the intended properties of these mock communities (**Supplementary Note 1**, **Supplementary Fig. 3, Supplementary Table 3)**. In brief, the DNA level representation of each species approximated its intended relative abundance in each mixture, with good agreement in *P. copri* levels between the arabinan and glucose conditions (**Supplementary Fig. 3a**). In contrast, *P. copri* transcripts constituted a higher proportion of metatranscriptomes in the glucose-conditioned mixtures compared to arabinan-conditioned mixtures, suggesting increased transcriptional output in glucose (**Supplementary Fig. 3b**). For initial method benchmarking without this confounder, we controlled for these changes by rarefying^71–73^ (sub-sampling without replacement) MTX counts from the mixtures containing glucose-conditioned *P. copri* to balance the species-level (*P. copri* and *E. coli*) transcriptional output between the conditions (see *Methods*). In this case, loss of available information due to rarefaction was modest because the range of differential representation was low (as opposed to multiple orders of magnitude). We confirmed that rarefaction resulted in even representation of *P. copri* transcripts in metatranscriptomes from both conditions while preserving gene-level quantification^74,75^ relative to each organism’s total transcriptome (**Supplementary Fig. 3d,h-j**). However, because transcription rate changes at the organism-level are a possibility in real-world datasets and may signify bacterial growth and death^39^, we also repeated our benchmarking using the original metatranscriptomes (**Supplementary Note 2**).

#### Defining true positive and true negative genes

To assess the sensitivity and specificity of different methods across comparisons with confounding low relative abundance, differential abundance, and non-colonization, we defined the ‘ground truth’ of differentially expressed genes by applying the widely used single organism RNA-seq method, DESeq2, to the 100% *P. copri* samples. Sets of true positive and true negative genes with strong evidence for respective presence or absence of differential expression were defined using threshold cutoffs for log fold-change and adjusted *P*-value (**Fig. 3b**, **Supplementary Table 4;** see *Methods*). Other benchmarked statistical methods produced true positive and true negative gene sets consistent with those defined by DESeq2, indicating that determination of ground truth was robust to tool choice (**Fig. 3c**). True positive rate, false positive rate, positive predictive value, and negative predictive value were then defined by the fractions of the ground truth gene sets which were inferred as differentially expressed by other tools (see *Methods*).

To confirm that the true positive genes identified statistically were also biologically plausible, we used homology-based annotations of carbohydrate active enzymes (CAZymes) and polysaccharide utilization loci^76^ in *P. copri* as well as domain-based annotations of predicted open-reading frames^77^. The most upregulated genes in each condition were two sets of genomically-adjacent, predicted glycoside hydrolase (GH) genes and susC/D pairs (**Fig. 3d**). The GHs upregulated in arabinan had predicted α-L-arabinofuranosidase and endo-arabinanase activities, while those upregulated in glucose encoded CAZymes with predicted α-glucosidase and α-amylomaltase activity (**Supplementary Table 4**). Additionally, growth on arabinan resulted in more modest but significant upregulation of genes involved in arabinose catabolism via the pentose phosphate pathway^78,79^, including a predicted ribulokinase, L-arabinose isomerase, L-ribulose-phosphate epimerase, ribose-5-phosphate isomerase B, and transketolase (**Fig. 3e; Supplementary Table 4**).

#### Benchmarking methods under confounding conditions

We used the defined mixtures of *P. copri* and *E. coli*, as well as the true positive and true negative gene sets defined above, to compare method sensitivity, specificity, and precision under various confounders (**Fig. 3f**, **Supplementary Fig. 4; Supplementary Table 5**). We employed receiver operating characteristic area under the curve (ROC AUC) as a summary metric of differential expression classification (**Fig. 3g-i**) in comparisons confounded by low relative abundance, differential abundance, and non-colonization. All methods demonstrated a gradual loss of sensitivity and differential expression classification in comparisons where low relative abundance led to insufficient transcriptome coverage (**Fig. 3f,g; Supplementary Fig. 4a**). None of the approaches for controlling for abundance effects (i.e., taxon-scaling or DNA covariates) were able to prevent the effects of loss of information due to lower sequencing coverage. Based on the average number of *P. copri* mapped transcript reads for the 10%, 1%, 0.1%, and 0.01% *P. copri* mixtures, for the sample size of six replicates per condition used here we recommend minimum sequencing efforts of approximately 10^6^, 10^5^, 10^4^, and 10^3^ reads successfully mapped to a genome of interest to recover approximately 80%, 50%, 20%, and 5% of its differentially expressed genes, respectively (**Fig. 3f**). These recommended depths are likely underestimates for human and animal studies where inter-sample variance is expected to be greater and statistical power lower compared to *in vitro* samples.

Confounding differential abundance was emulated by performing all pairwise comparisons between *P. copri* mixtures in arabinan and glucose (**Fig. 3h; Supplementary Fig. 4b**). As expected, community-scaled DESeq2 (CSS) without abundance information, or taxon-scaling, exhibited an extremely high false positive rate (up to 98%) leading to poor classification performance. Both DNA-normalized and taxon-scaled DESeq2 maintained sensitivity and controlled false positives except at extreme magnitudes of differential abundance that are unlikely in real datasets (i.e., 100% vs. 0.01%). Taxon-scaled MPRAnalyze had lower sensitivity, specificity, precision, and AUC compared to DNA-normalized and taxon-scaled DESeq2. MTXmodel, despite controlling for DNA abundance information and demonstrating high sensitivity on the simulated datasets, failed to recover differentially expressed genes in the presence of confounding differential abundance.

Zero-inflation from confounding low prevalence was emulated by including either zero, two, six, or eighteen 0% *P. copri* samples in each group to achieve prevalences of 100, 75, 50, and 25% (**Fig. 3i; Supplementary Fig. 4c**). All DESeq2 approaches were susceptible to this confounder and demonstrated decreasing differential expression classification with decreasing prevalence. In contrast, MPRAnalyze and MTXmodel, both of which utilize abundance data and filter out samples with zero DNA counts, were robust to decreased prevalence and maintained all performance metrics. We concluded that DESeq2 with either DNA covariates or taxon-scaling demonstrated the highest sensitivity, specificity, precision, and ROC AUC of all tested methods if an organism had low relative or differential abundance, but not when its prevalence was low.

#### Additional validation benchmarking analyses

To ensure that using DESeq2 CSS to define true positive and true negative genes was not a source of methodological bias favoring performance of DESeq2-based methods, we defined separate ‘consensus’ benchmarking gene sets (571 true positives and 1,304 true negatives) taken as the union across the respective sets defined by each tested method when applied to the monoculture samples (**Supplementary Fig. 5a**). We then separately calculated performance metrics using these gene sets. Sensitivity, specificity, precision, and AUC were slightly lower across methods because no individual method completely captured these consensus sets (**Supplementary Fig. 5b-e**). However, the robustness and susceptibility of methods to low relative abundance, differential abundance, and varying prevalence were unchanged (**Supplementary Fig. 5e-g**).

We separately performed benchmarking using the original (non-rarefied) counts profiles, where *P. copri* had demonstrated higher transcriptional output in the glucose condition that was not observable in the monoculture context due to the relative nature of sequencing counts (**Fig. 1b,c**). Although this prevents consistent identification of ‘ground truth’ gene upregulation between monocultures and mixtures, we leveraged this feature of the metatranscriptomes to determine whether methods could recover upregulated genes in both conditions despite global transcription rate changes, as is possible in single-organism analysis. Given the larger fraction of RNA reads from *P. copri* in glucose, we additionally hypothesized that the compositional nature of sequencing data (i.e., the non-independence of features measured as relative quantities^80–81^) would lead to lower counts per gene for *E. coli*, resulting in false positives despite the lack of differences in *E. coli* growth conditions. Taxon-scaling robustly controlled false positives due to compositional effects on *E. coli* metatranscriptomic representation and successfully inferred upregulation in both the arabinan and glucose conditions. Other methods were prone to false positives due to compositionality and failed to infer genes upregulated in arabinan (**Supplementary Note 2, Supplementary Fig. 6, Supplementary Tables 6,7**).

We next determined whether the observed discrepancy in the performance of methods in simulated datasets and mock communities was due to other confounding differences between the datasets. To do so, we used the same previously published^50^ metagenomic and metatranscriptomic simulation framework to generate additional simulated datasets with (i) equivalent sample sizes of 6 per condition but maintaining complexity of 100 species and (ii) both equivalent sample size and equivalent complexity of two species with the relative abundances used in our mock communities (**Supplementary Fig. 7a,b;** *Methods*). We quantified sensitivity, specificity, precision, and ROC AUC on these new simulated data. In both the complex (100 species) and two species synthetic communities, MTXmodel exhibited the highest sensitivity and best differential expression classification with the highest AUC (**Supplementary Fig. 7c,d**). The performance difference between MTXmodel and taxon-scaled DESeq2 was much more pronounced in the two-species communities, where MTXmodel demonstrated near-perfect sensitivity and AUC at high simulated relative abundances (**Supplementary Fig. 7d**). These results indicate the relative performance of methods in synthetic datasets does not depend on sample size or community complexity, and that the simulation framework likely encodes assumptions (e.g. direct scaling of RNA expression by DNA abundance) that inflate the performance of MTXmodel.

#### Recommendations for metatranscriptomic differential expression

Based on these results, we conclude that DESeq2 with taxon-scaling adequately controls false positives due to differential abundance, transcription rate changes, and compositional effects, and demonstrated sensitivity that outperformed other methods except in the presence of low prevalence. These results contrast with method performance on simulated data, highlighting the importance of benchmarking under realistic conditions. We recommend taxon-scaled DESeq2 for uses where inter-sample heterogeneity is low and all samples are colonized with the organism of interest, such as *in vitro* or gnotobiotic animal experiments (see below). If prevalence is incomplete, use of abundance information by MTXmodel or taxon-scaled MPRAnalyze maintains sensitivity, with the choice between the two representing a sensitivity-specificity tradeoff. Additionally, for the sample size of six replicates used here, we recommend minimum sequencing efforts of approximately 10^6^, 10^5^, 10^4^, and 10^3^ reads mapping to an organism to respectively provide sufficient coverage for inference of 80%, 50%, 20%, and 5% of its differentially expressed genes.

### Differential expression in the intestinal metatranscriptomes of gnotobiotic mice

Based on performance differences of methods using simulated and *in vitro* benchmarking datasets, we aimed to test whether they produced consistent results and supported the same conclusions when applied to real biological datasets. Similarly, we sought to demonstrate that careful consideration of differential expression methodology can empower new biological insights.

We used MTXmodel, which quantifies changes in RNA representation relative to DNA representation, community-scaled DESeq2, which quantifies changes in RNA representation, and taxon-scaled DESeq2, which quantifies changes in relative RNA representation within each organism, to analyze metagenomic and metatranscriptomic datasets from a previously published gnotobiotic mouse study^70^. In this experiment, one group of animals was colonized with a defined consortium of 19 cultured, genome-sequenced human gut bacterial strains including *P. copri* and fed a diet rich in plant glycans^65,69^. Another group was colonized with the same community but without *P. copri* and fed the same diet (**Fig. 4a**). Comparison of the two groups of gnotobiotic mice previously disclosed that *P. copri* colonization was associated with increased fitness (absolute abundance) of several organisms^70^, introducing confounding differential abundance for metatranscriptomic analysis. We used these two groups to assess transcriptional responses of other organisms to the presence of *P. copri* in communities sampled at the time of euthanasia from the cecum, and to test whether *P. copri*-mediated glycan degradation was associated with altered nutrient utilization by other microbial species.

**Figure 4.**
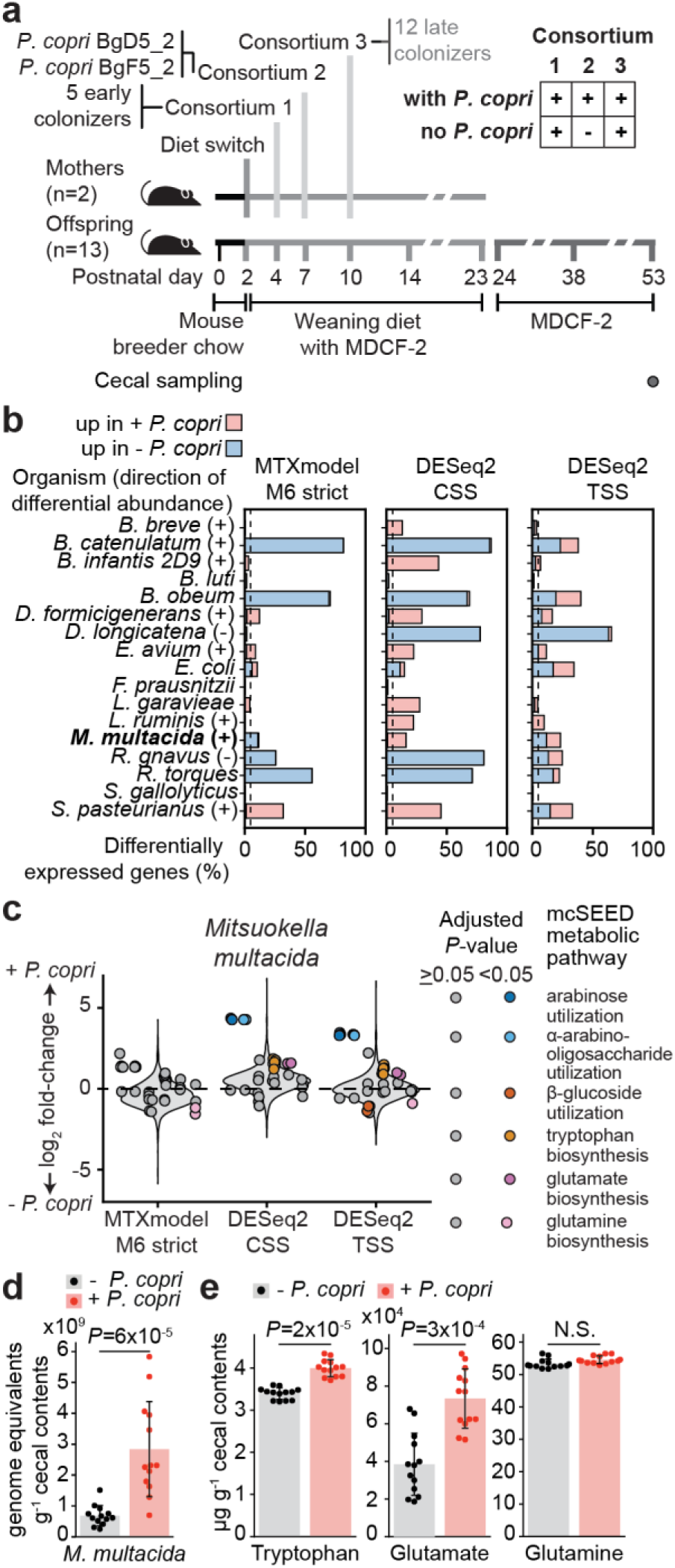
Metatranscriptomic differential expression nominates cross-feeding interactions from a gnotobiotic mouse model. **(a)** Experimental design testing effects of *P. copri* colonization in a defined model of the human gut microbiota, adapted from a previous publication^70^. (**b**) Percentage of differentially expressed genes from each organism inferred by each method. Taxon names are annotated to indicate when an organism’s abundance was significantly increased in the presence (+) or absence (-) of *P. copri*. Colors indicate whether differentially expressed genes were inferred as upregulated (log fold-change > 0) in the + *P. copri* or – *P. copri* group by MTXmodel, DESeq2 with community-scaling, and DESeq2 with taxon-scaling. The dotted vertical line shows the alpha threshold of 0.05. (**c**) Differential expression results for genes predicted to be involved in mcSEED metabolic pathways involved in the utilization of arabinose and alpha-arabinosides, as well as biosynthesis of glutamate, glutamine, and tryptophan for *Mitsuokella multacida* as reported by the three methods using an adjusted *P*-value threshold of 0.05. Violin plots represent the distribution of log_2_ fold-change across all genes and points represent differential expression test results for individual genes from select metabolic pathways, horizontally grouped by pathway and colored if they were significantly differentially expressed according to a given method. The dashed line indicates no change in expression (a log_2_ fold-change value of 0). (**d**) Absolute abundance of *M. multacida* estimated relative to spike-in bacteria. (**e**) Cecal tryptophan, glutamate, and glutamine levels quantified by targeted mass-spectrometry. Mean ± standard deviation values are shown for panels d and e. *P*-values were calculated by the specified differential expression methods in panel c, and a two-sided Mann-Whitney U test in d and e.

Taxon-scaled DESeq2 and MTXmodel identified a similar mean fraction of differentially expressed genes per organism (19.7% and 18.8%, respectively) that was lower than community-scaled DESeq2 (36.6%), though all of these were markedly higher than the null hypothesis expectation of 5% (**Fig. 4b**). However, MTXmodel and community-scaled DESeq2 both produced results where differentially expressed genes within each organism were largely in the same direction as differential abundance of organisms (**Fig. 4b; Supplementary Table 8a**). In contrast, DESeq2 with taxon-scaling more evenly inferred upregulated genes in both the *P. copri* colonized and uncolonized groups.

Previous transcriptional and mass-spectrometry data from these animals indicated that *P. copri* prioritized expression of its arabinan-targeting polysaccharide utilization locus, and that *P. copri* colonization was associated with increased degradation of arabinose-containing glycans in the cecum^70^. Given that this locus encodes predicted extracellular arabinofuranosidases, we hypothesized that *P. copri* might liberate arabinose that could cross-feed other organisms whose genomes encode fewer enzymes involved in polysaccharide degradation. Consistent with this hypothesis, we identified upregulated genes involved in free arabinose utilization in three of the four members of the defined community with predicted capabilities to utilize it. Of these arabinose-utilizing organisms, we focused on gene regulation in *Mitsuokella multacida* after observing that inferred results differed among the methods applied (**Fig. 4c; Supplementary Table 8b**). We identified significant upregulation of *M. multacida* pentose phosphate pathway genes involved in utilization of arabinose and arabino-oligosaccharides only using DESeq2, regardless of the scaling method applied. However, significant upregulation of genes in the ‘no *P. copri* group’, such as those in *ß*-glucoside utilization and glutamine biosynthesis, was only apparent when using taxon-scaling (**Fig. 4b,c**). *M. multacida* also exhibited upregulation of genes involved in glutamate and tryptophan synthesis in DESeq2-based analyses. MTXmodel did not infer significant upregulation for genes from any of these pathways.

*P. copri* colonization was associated with a four-fold increase in the absolute abundance of *Mitsuokella multacida*, suggesting that these changes in carbohydrate utilization responses may increase its fitness (**Fig. 4d**). Targeted mass spectrometry disclosed that *P. copri* colonization was associated with increased cecal levels of glutamate and tryptophan, correlating with upregulation of their biosynthetic pathways by *M. multacida* (**Fig. 4e**). Based on these results, we hypothesized that *M. multacida* fitness, utilization of arabinose, and amino acid biosynthesis partially depend on *P. copri* liberation of arabinan mono- and oligosaccharide components.

We subsequently validated this potential cross-feeding relationship using *in vitro* mono- and cocultures of *P. copri* and *M. multacida* (**Supplementary Note 3, Supplementary Fig. 8, Supplementary Table 9-13**). Using optical density measurements and quantitative PCR (qPCR) assays for each organism, we found that although *M. multacida* could grow alone in a defined medium^82^ with either arabinose or glucose as the sole carbon source, it required the presence of *P. copri* to grow with arabinan as the sole carbon source. We confirmed that *M. multacida* upregulated both arabinose utilization as well as tryptophan and glutamate biosynthesis pathways in response to free arabinose availability and *P. copri*-mediated arabinan cross-feeding, just as it had in response to *P. copri* colonization in the gnotobiotic animal model. Successful differential expression inference of these genes from co-cultures was possible using taxon-scaled DESeq2 but not MTXmodel. Targeted mass spectrometry confirmed that *Mitsuokella* upregulation of these biosynthesis pathways was associated with low availability of tryptophan and increased availability of glutamate in the conditioned media. Based on these results, we concluded that taxon-scaled DESeq2 was able to nominate cross-feeding interactions and downstream amino acid metabolic responses from our gnotobiotic animal studies which could then be validated *in vitro*.

### Enhancing differential expression inference in human studies

Taxon-specific scaling demonstrated robustness to all confounders except for its low sensitivity with low organism prevalence. To determine whether this limitation could be overcome in human studies where few bacterial strains are present across all participants, we turned to paired metagenomes and metatranscriptomes generated from fecal samples collected from participants in a clinical study comparing therapeutic foods for childhood undernutrition^65,70^. We used differential expression to identify transcriptional responses to a microbiota directed complementary food formulation (MDCF-2), by comparing transcriptomes before and at the end of treatment. We reasoned that analogous to the sample-level zero-filtering implemented by MTXmodel and MPRAnalyze, we could identify and exclude samples with insufficient sequencing coverage of a given organism.

We first defined two metrics, genome-level depth (*A*) and gene-level detection (*D*), which respectively represent a genome’s total counts across its encoded genes and the fraction of its genes with non-zero counts in each sample (*Methods*). We initially quantified these metrics in our mock community samples, where high detection in the pure *P. copri* samples allowed us to definitively ascribe decreases in detection at lower genome-level depths to technical causes (insufficient sequencing coverage) and not to actual lack of expression (**Fig. 5a**). The detection of genes in RNA transcripts better approximated the sensitivity of differential expression than the detection of genes in community DNA; we surmised this was likely due to uneven coverage of the transcriptome causing loss of detection for lowly-expressed genes at higher genome-level depths of coverage in community RNA compared to community DNA. Metagenome-assembled genomes (MAGs) from the human study exhibited a similar relationship between depth and detection, albeit with slightly lower detection and higher variance for equivalent genome-level depth of coverage (**Fig. 5b**).

**Figure 5.**
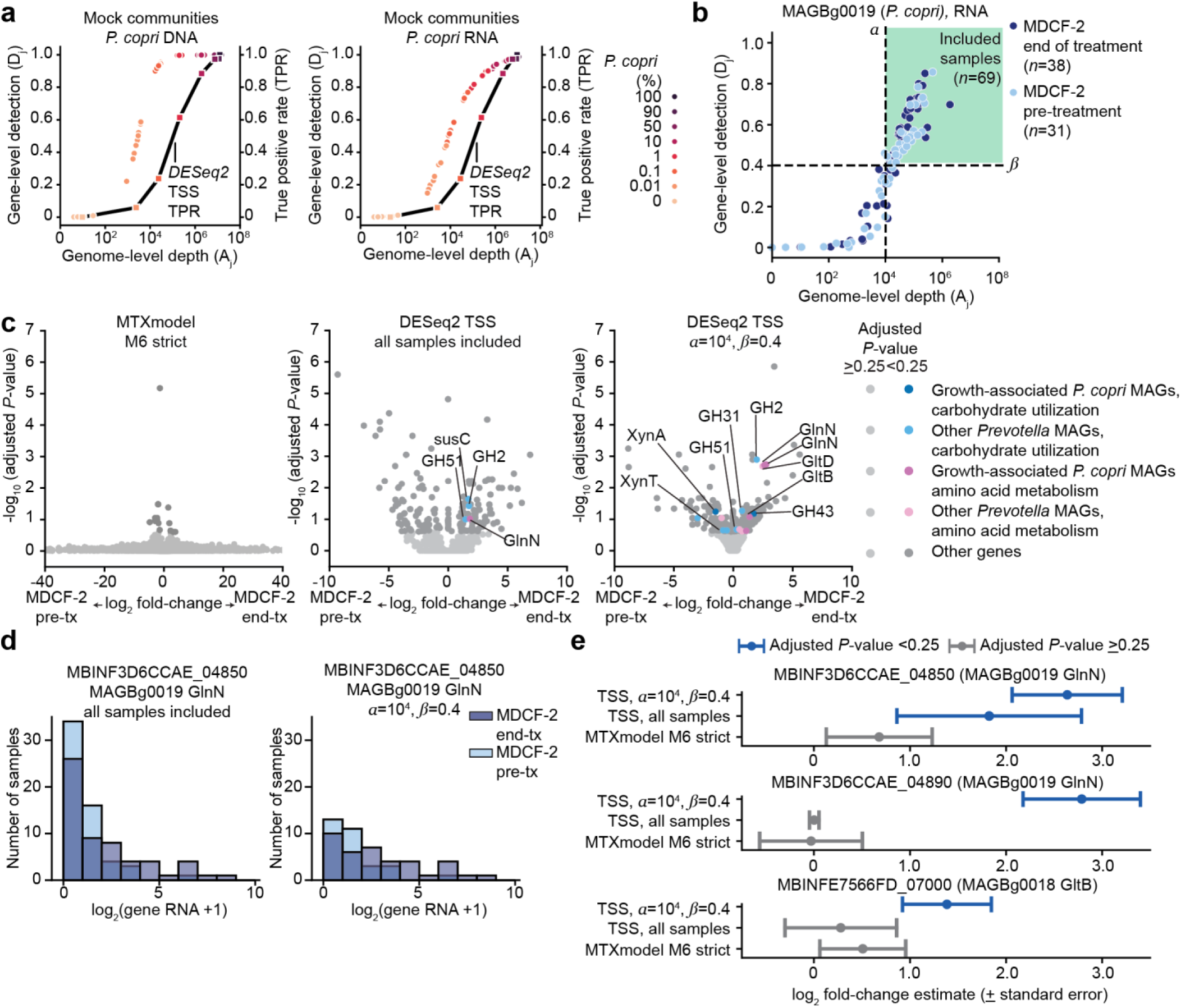
Depth and detection filtering of samples increases differential expression inference in a human study. **(a)** Genome-level depth (A_j_) and gene-level detection (D_j_; see *Methods*) calculated using DNA counts or RNA counts in mock community samples described in Fig. 3, plotted with the sensitivity of taxon-scaled DESeq2 for samples at each mixture ratio. **(b)** Depth and detection plotted for a single metagenome-assembled genome (MAG) for *P. copri* whose abundance had previously been associated with increased ponderal growth (weight-for- height Z score) in study participants^70^. The depth and detection thresholds used in subsequent panels are plotted with vertical and horizontal dashed black lines, respectively, and samples included under these thresholds are shown in the upper right green square. **(c)** All differential expression results using either MTXmodel (M6 strict) or taxon-scaled DESeq2 including all samples or using depth and detection filtering with *α*=10^4^ and *ß*=0.4. Genes from MAGs assigned to the genus *Prevotella* or from two ponderal growth-associated *P. copri* strains (MAGs)^70^ are shown in light and dark colored sets of points. Genes with functional annotations in carbohydrate utilization pathways are shown in blue and genes with functional annotations in amino acid metabolic pathways are shown in pink. Specific genes involved in utilization of arabino-oligosaccharides, beta-glucosides, or xylo-oligosaccharides as well as biosynthesis of glutamate or glutamine have their names annotated. Statistical significance was defined using a cutoff of 0.25 for *P*-values adjusted across all genes from all genomes with sufficient replicates for analysis (up to 1,929,056 genes with no sample filtering). **(d)** Representative distributions of gene-level counts for the *P. copri* glutamine synthetase gene that was differentially expressed both when considering all samples and when using the depth and detection filtering in panel c. **(e)** Log_2_ fold-change and standard error estimates (points and lines respectively) for glutamine and glutamate biosynthesis genes from growth-associated *P. copri* MAGs shown in panel c, using MTXmodel or taxon-scaled DESeq2 with all samples or depth and detection filtering. Parameter estimates are colored if they met statistical significance. Abbreviations: TSS, taxon-specific scaling; pre-tx, pre-treatment; end-tx, end of treatment; GH51, glycoside hydrolase family 51 (exo-α-1,2-arabinofuranosidase); susC, TonB-dependent transporter; GH2, glycoside hydrolase famly 2 (beta-galactosidase); GlnN, glutamine synthetase; XynT, xyloside transporter; XynA, endo-1,4-beta-xylanase; GH31, glycoside hydrolase family 31 (alpha-xylosidase); GH43, glycoside hydrolase family 43 (endo-α**-**1,5-arabinanase); GltB, glutamate synthase large chain; GltD, glutamate synthase small chain.

To reduce zero-inflated counts from samples where gene-level detection was low for a given genome-level depth, we opted to exclude samples by considering both depth and detection; we hypothesized that this would enhance differential expression inference in the human study. We performed a parameter sweep of thresholds to select cutoffs for exclusion of samples from a given genome’s differential expression analysis (**Supplementary Fig. 9; Supplementary Table 14)**. Excluding samples represents a tradeoff between decreasing statistical power and generalizability across individuals while increasing confidence in quantification by decreasing zero-inflation in the remaining analyzed samples. Excluding samples decreased the number of genomes with sufficient samples for statistical testing, but increased inference of differentially expressed genes (**Supplementary Fig. 9a-d**).

We then sought to identify genes for which differential expression inference was robust to different degrees of sample exclusion. We defined a set of genes that were differentially expressed across 12 (20%) of the threshold pairs, encoded by higher abundance organisms, and supported by a greater number of included samples (*Methods*; **Supplementary Fig. 9e-g; Supplementary Table 15**). We selected depth and detection thresholds of 10^4^ and 0.4, respectively, because they exhibited high recovery of this set and maximized its fraction of all differentially expressed genes (**Supplementary Fig. 9h,i**). We applied these thresholds to our mock communities and found that while they decreased inference at low relative abundances, they increased robustness to confounding low prevalence (**Supplementary Fig. 9j-l**). We subsequently used these thresholds for comparisons of exclusion and inclusion of low information samples in the human study.

Compared to including all samples, taxon-scaled DESeq2 with these depth and detection thresholds increased differential expression inference approximately two-fold and reduced the spread of fold-change estimates for non-significant genes (**Fig. 5c; Supplementary Fig. 9c; Supplementary Table 16**). MDCF-2 treatment was associated with utilization of arabino-oligosaccharides via a GH51 exo-α-1,2-arabinofuranosidase and *ß*-galactosides via a GH2 *ß*-galactosidase in both analyses (**Fig. 5c**). Sample filtering also recovered upregulation of synthase and synthetase genes involved in glutamate and glutamine biosynthesis and a GH43 endo-α**-**1,5-arabinanase in a set of *P. copri* genomes whose abundances had been previously positively associated with weight gain in the study participants^70^ (**Fig. 5c**). Depth and detection filtering decreased the degree of zero-inflation of gene-level counts for these genes by excluding cases where zero counts could likely be attributed to insufficient sequencing depth; this increased the magnitude and decreased the standard error for their estimated log_2_ fold-changes (**Fig. 5d,e**).

We conclude that differential expression inference with taxon-scaling can be increased for human cohort studies by identifying and excluding samples that have insufficient depth of RNA coverage and detection of genes for each genome. We propose starting point depth and detection thresholds of at least 10^4^ RNA counts for a genome with 40% of its genes detected, although this approach can be generalized to study the transcriptional responses of lower or higher abundance organisms by decreasing or increasing these thresholds.

## DISCUSSION

Despite the accelerated generation of metagenomic, metatranscriptomic, and mass-spectrometric datasets intended to characterize functional activities of bacteria within microbial communities, deciphering the mechanisms underlying microbial-microbial and host-microbial interactions remains challenging. In this study, we present a framework for evaluating the accuracy of inferring differential gene expression for individual organisms within microbial communities. Rather than the ‘top-down’ approach adapted by studies using data simulated to represent highly diverse communities, we employed a ‘bottom-up’ approach using simple, real bacterial samples to test the effects of modular increases in complexity and confoundedness on method performance. We demonstrate that current commonly used methods are limited in their ability to recover differentially expressed genes and control false positives under confounding low relative abundance, differential abundance, variable prevalence, global alterations in transcription rate, and compositional effects. We show that existing methods differ in their ability to infer experimentally validated cross-feeding interactions in gnotobiotic mice colonized with a defined consortium of cultured genome-sequenced human gut bacterial strains. We leverage genome-level depth and gene-level detection of genomes to reduce zero-inflation and overcome confounding low prevalence to enhance differential expression inference from a human clinical study. Together, these analyses yield recommendations for sequencing effort and choice of analytical method as well as improvements to existing approaches for microbial community differential expression based on properties such as study type, community composition, heterogeneity, differential abundance, and prevalence.

This study assumes a goal of inferring symmetric upregulation of genes that is robust to changes in organism-level transcription rate, as would be obtained in single-organism transcriptomics. While this is informative for interpreting transcriptional responses of individual microbes to environmental perturbations, there are cases where the aggregate community-level functions are of interest. As such, inference of ‘community-scaled’ and relative-to-organismal-abundance transcription (e.g., MTXmodel) are also useful, although they underperformed by the metrics in our benchmarking study as well as in animal and cross-feeding experiments.

Our benchmarking strategy was based on analyses of low complexity mock communities which are easier to reproducibly manipulate^52^. While low-complexity communities exist in some body habitats (e.g. vaginal and skin microbiota^83–85^), application of metatranscriptomic sequencing to the human gut microbiota requires methods that perform in the presence of hundreds of distinct and often genomically-similar organisms, such as in the case of >100-member synthetic defined communities that enable mechanistic interrogation of metabolic niches^86,87^. Because differential expression inference of individual organisms does not directly depend on community complexity, our benchmarking approach used complete and uncontaminated reference genomes to enable reliable quantification of microbial genes of interest. Nonetheless, when analyzing complex human communities where reliable genome assembly and gene quantification present challenges not encountered in our mock community framework, taxon-scaling recovered transcriptional responses of individual microbes and demonstrated improved inference when leveraging depth and detection filtering. We anticipate that with continued investment in metagenomic methods designed for use in complex, intact communities, the capacity to study microbial community gene expression in human cohorts will continue to improve.

We recommend taxon-scaled DESeq2 for metatranscriptomic analyses where the prevalence of organisms of interest is high. We demonstrate that sample exclusion using genomic metrics of depth and detection can mitigate the confounding effects of low prevalence of taxa in human studies. However, zero-inflation of metatranscriptomic data from human studies remains a significant challenge; as such, new statistical methods designed explicitly for metatranscriptomic differential expression analyses and tested using the framework described here are needed. Such approaches could leverage the genome-level structure^38^ within microbial communities as we have here, or apply existing empirical Bayes approaches for parameter estimate moderation^45^ and/or utilize the coregulation of genes within bacterial operons to improve differential expression inference. These methods must handle the inherent compositionality, biological and technical zero-inflation, and abundance-dependence of metatranscriptomic sequencing data. Such advances will enable the next generation of studies where inference of the key functional activities executed by specific organisms in their community context should yield new insights into the mechanisms by which the human gut microbiome influences health and disease.

## Acknowledgements

We thank M. Meier for his essential role in generating metagenomic and metatranscriptomic libraries and sequencing datasets, D. O’Donnell and M. Karlsson for their assistance with mouse husbandry, and J. Guruge for help with culturing and maintaining bacterial strains.

## Contributions

E.M.L., N.P.M., M.C.H., B.A.C., and J.I.G. designed the mock community and cross-feeding *in vitro* experiments. E.M.L., H.-W.C., and J.I.G. designed the gnotobiotic mouse experiments. All bacterial and animal experiments were performed by E.M.L. Metagenomic and metatranscriptomic datasets were processed and analyzed by E.M.L and M.C.H. with workflows designed by E.M.L. and M.C.H., with assistance from N.P.M. Mass spectrometry-based metabolomic studies were conducted by J.C. E.M.L., N.P.M., B.A.C., and J.I.G. wrote this manuscript with invaluable assistance from co-authors.

## Data availability

The datasets generated by shotgun sequencing of DNA and RNA extracted from mock community samples and cross-feeding coculture experiments have been deposited in the National Center for Biotechnology Information Sequence Read Archive (SRA) under study accession number PRJNA1293448. DNA and RNA sequencing data from previously published gnotobiotic mouse studies are available in SRA under study accession number PRJNA1067830. Annotated genomes for the strains of *P. copri*, *E. coli*, and *M. multacida* used in this study have been deposited in the GenBank Genome collection. Processed counts datasets are available in Supplementary Information and upon request from the corresponding author.

## Code Availability

Python code for processing sequencing reads quantification, differential expression results, and the calculation and visualization of benchmarking performance metrics, as well as R scripts for running the various differential expression methods are available on GitHub at https://github.com/evanmlee/MTX_utils and https://github.com/evanmlee/MTX_DE. All other software used were from publicly available sources.

## METHODS

### Quantitative Data Analysis

Unless otherwise specified, metagenomic and metatranscriptomic counts data, differential expression results, qPCR and growth data from bacterial cultures, and mass-spectrometry data were analyzed in Python using numpy^88^, scipy^89^, and pandas^90^, and visualized with seaborn^91^ and matplotlib^92^.

### Statistics and reproducibility

Sample sizes for bacterial culture experiments were not pre-determined by statistical methods but are close to those in previous publications^70,82^. For differential expression and other statistical testing, a significance threshold (alpha) of 0.05 was used unless otherwise stated. *P*-values were adjusted using the Benjamini Hochberg

### Differential expression method benchmarking using simulated datasets

Six simulated datasets from Zhang et al.^50^ were used for method benchmarking. In brief, these datasets each simulated both metagenomic (MGX) and metatranscriptomic (MTX) counts for 100 samples based on the abundance and prevalence of the 100 most-abundant organisms in data from the Human Microbiome Project^62^. For each organism, MGX and MTX counts were sampled from log-normal distributions scaled by empirical abundance for the fraction of samples simulated as colonized based on empirical prevalence data, and assuming no effect on simulated counts from variations in gene length or variation in gene encoding frequencies within or across taxa. A subset of 10% of genes were simulated as differentially expressed in either the positive or negative direction by reassigning sampled MTX expression counts to induce rank correlation with sample metadata (‘phenotype’).

Differential expression methods (described below) were applied to these datasets and performance was assessed using the true positive rate (TPR) and false positive rate (FPR). TPR was quantified as the fraction of genes which were simulated as differentially expressed (‘spiked’) for which tests yielded a significant result (adjusted *P*-value < 0.05) in the correct direction (increased vs. decreased expression). For consistency with reporting in Zhang et al., FPR was quantified using nominal (i.e., not adjusted) *P*-values and was calculated as the fraction of genes which were not simulated as differentially expressed that were falsely tested as differentially expressed (nominal *P*-value < 0.05).

Positive predictive value (PPV) and negative predictive value (NPV) were also estimated from differential expression results. PPV was defined as the fraction of positive test results, that is both true and false positives defined by an adjusted *P*-value < 0.05, which corresponded to genes that were simulated as differentially expressed. NPV was defined as the fraction of negative test results, that is both true and false negatives defined by a nominal *P*-value > 0.05, corresponding to genes which were not simulated as differentially expressed. Both these metrics depend on the prior distributions of ground truth differentially expressed vs. non-differentially expressed genes, which were approximately 10% and 90% of all genes, respectively. Positive predictive value and negative predictive value were defined as 0 in edge cases where there were respectively no positive test results or no negative test results.

For receiver operating characteristic area under the curve (ROC AUC) quantification in **Fig. 2g**, additional modifications were made for compatibility with the scikitlearn ROC implementation^93^. Both TPR and FPR were both calculated using adjusted *P*-values since only one y_score metric is supported. In cases where differential expression testing was skipped for genes or genomes with insufficient abundance, *P*-values of 1 (no differential expression) were imputed to prevent inflation of method performance. This is most apparent for DESeq2 CSS DNA, which exhibited high precision for its predictions but skipped testing for large fractions (45-70%) of the datasets (**Fig. 2e**; **Supplementary Table 1b**), and in the ‘true-combo-bug-exp’ dataset where ‘strict’ MTXmodel zero-filtering resulted in skipped testing for 30% of genes. Similarly, because DESeq2 performs more manual manipulations on adjusted *P*-values by setting padj to NA for genes meeting certain criteria (all 0 counts, outlier counts, or low mean expression), this would artificially increase DESeq2’s area under the curve by filtering out many features with low expression due to low underlying microbial abundance. The number of genes with numeric (non-NA) *P*-values are listed for each simulated dataset and method in **Supplementary Table 1b**. Because genes with low expression or detection often met DESeq2’s criteria for missing *P*-values and similarly represent cases where inference of differential expression is challenging, these *P*-value manipulations would otherwise inflate estimation of TPR and FPR for DESeq2 in the AUC analysis. For more details on criteria for missing *P*-value entries, see the section ‘*More information on results columns*’ in the DESeq2 user vignette^45^.

For analysis of simulated datasets binned by underlying organism relative abundance, genes from all 100 organisms were sorted into deciles of 10 organisms by ranking each organism by its median DNA abundance in samples where it was present (i.e., not accounting for prevalence). Sensitivity, specificity, and precision were then calculated for the genes from each decile as before.

#### DESeq2

DESeq2^45^ is a widely-used method for differential expression analysis which was developed for single-organism RNA-seq. It uses negative binomial generalized linear models with a logarithmic link to estimate effects of covariates on expression. It employs Empirical Bayesian shrinkage of parameter estimates to moderate overdispersion and fold-change estimation of gene-wise estimates towards common trends for all genes. DESeq2 was run with ‘poscounts’ library size factor calculation to accommodate zero-inflated counts in samples with low abundance. By default, DESeq2 estimation of dispersions includes gene-level dispersion estimation and Empirical Bayes dispersion shrinkage towards a common trend. Fitting of a common trend fails if dispersion estimates are extremely small, such as for *P. copri* genes in the 0% *P. copri* samples, and in these cases, initial gene-level dispersion estimates were used without shrinkage. Log fold-change shrinkage was also applied with the function lfcShrink using priors from ashr^94^.

In the single-organism context that DESeq2 was developed for, the size factors used to control for variations in sequencing effort are calculated from the profile of all gene counts. For metatranscriptomics of multiple-member microbial communities, this is analogous to scaling at the community-level (community-sum-scaling). DESeq2 was also separately run implementing taxon-specific scaling by estimating size factors for genes from each individual genome separately^49^. *P*-values were separately corrected for multiple tests across genes from all genomes. This modified approach is referred to as taxon-scaled DESeq2, or DESeq2 TSS. Similarly, because DESeq2 does not support gene-level covariates, gene-level DNA abundances (normalized by row-wise geometric means, as recommended by the developers in the DESeq2 user’s vignette^45^) were optionally provided as a normalization matrix when estimating size factors such that size factors account for both RNA sequencing effort and DNA abundance. This approach is referred to as DESeq2 CSS DNA and DESeq2 TSS DNA when used with community-scaling or taxon-scaling.

#### MTXmodel

MTXmodel^50^ was developed specifically to account for confounding abundance-dependence in MTX data and has been widely-applied to environmental and gut microbial communities^54,55^. MTXmodel differential expression best practices have since been incorporated into MaAsLin3^28^, another widely-used tool for identifying associations between microbial features and sample covariates. MTXmodel assumes log-normal distributions and fits Gaussian generalized linear models to log-transformed RNA counts.

MTXmodel was applied using gene-level DNA and RNA counts for all datasets with no minimum prevalence, either zero-filtering settings of ‘lenient’ (removal of genes with all 0 counts in either DNA or RNA) or ‘strict’ (removal of samples for a given gene’s model if either DNA or RNA counts were 0). As initially described by Zhang et al.^50^, DNA abundance information was either not provided (M1), summed into taxon-level abundance covariates (M5), or used as gene-level DNA counts (M6). All other options used default values for the package.

#### MPRAnalyze

MPRAnalyze^63^ was developed for analysis of massively-parallel reporter assay data, where plasmid libraries incorporating different barcoded putative regulatory elements drive expression of a reporter gene to compare their relative effects on transcription rate. Because plasmids in the library are not transfected uniformly, normalizing to DNA abundance of each construct represents a similar analytical challenge as is seen in MTX differential expression. MPRAnalyze applies linked generalized linear models to separately model the effects of covariates on the DNA and RNA data, with the final reported ‘relative transcription rate’ having confounding effects from differential abundance of DNA counts removed. MPRAnalyze was run using gene-level DNA and RNA counts and treating the entire profile as a single barcoded library. Total sum library size correction was applied to both DNA and RNA libraries and differential expression tests were performed with the likelihood ratio test (testLrt). Library size factors were either applied at the community level (CSS) or by employing taxon-specific-scaling (TSS).

### Bacterial genome sequencing and annotation

All cultured isolates were obtained from fecal samples or are DSM catalog isolates, as described previously^70^. Bacterial genomes were sequenced, assembled, and putative open reading frames were annotated by comparison to both mcSEED metabolic pathways as well as predicted polysaccharide utilization loci (PULs)^70,76,77^.

### Bacterial culture for preparation of mock communities

Monocultures of each isolate were grown from frozen stocks overnight at 37 °C in Wilkins-Chalgren Anaerobe Broth [WC broth] (Oxoid; CM0643) in a Coy Chamber under anaerobic conditions (atmosphere: 75% N2, 20% CO2, and 5% H2) without shaking. Monocultures were streaked onto 10% horse blood agar plates and grown for 2 d at 37 °C to obtain single colonies. Individual colonies were picked into WC broth and grown for 24 h at 37 °C. A 1.5 mL aliquot was taken from this subculture for genomic DNA extraction and confirmation of a strain’s identity by 16S rDNA PCR and Sanger sequencing of the resulting amplicons. The WC broth subculture was then diluted 1:100 and grown for 24 h in *P. copri* defined medium [PCDM]^72^ with either 1% arabinan (Megazyme; P-ARAB) or 1% (w/v) glucose (Sigma; G8270) as the sole carbohydrate source for *P. copri* cultures and all other organisms, respectively. These acclimatization cultures in PCDM were standardized by OD_600_ across organisms and then used as the inoculum for the experiments described below.

To obtain estimates of colony-forming units (CFUs) for each organism at mid-log growth, each organism was acclimatized in PCDM as above and grown in 5 mL cultures of PCDM with 1% arabinan or 1% glucose. These cultures were prepared anaerobically in crimp-sealed culture tubes (Chemglass; CLS-4209) and brought out of the Coy chamber for growth at 37 °C and serial optical density measurements at 600 nm (Spectronic 20D+ spectrophotometer; Fisher Scientific; 14-385-130). Once cultures reached mid-logarithmic growth in an OD range of 0.45–0.6, they were returned to the Coy chamber and a 12-point 1:10 serial dilution was performed. A 10 µL volume for each point of this serial dilution was spotted onto a blood agar plate and spread with an inoculating loop. Colonies were counted 24–48 h later for each spot to obtain an estimate of CFUs/mL for each organism/media combination at its final measured OD.

To prepare mock community mixtures as described in **Fig. 3**, each organism was acclimatized in PCDM as described above, then subcultured in PCDM with the appropriate carbohydrate source as either one 10 mL culture for *P. copri* or two 15 mL paired cultures (30 mL total) for *E. coli* per biological replicate (n=6). For each biological replicate set of cultures of each organism, the 8 mixture ratios specified in **Fig. 3** were prepared as technical replicates by using the same source cultures as inputs for each ratio. Cultures were prepared in crimp-sealed tubes as described above and grown at 37 °C under anaerobic conditions to mid-logarithmic phase with final ODs ranging from 0.45 to 0.6. Cultures were then unsealed in the Coy chamber, resuspended, and transferred to 15 mL or 50 mL conical tubes, for *P. copri* and *E. coli* cultures respectively, before centrifuging (4 °C; 3,150 × g; 10 min). Conditioned medium was decanted, and a 2 mL aliquot was saved for mass spectrometry. Pellets were then resuspended in 2 mL of RNAlater per 10 mL of culture (Thermo Fisher; AM7021). RNAlater resuspensions were stored at 4 °C per the manufacturer’s recommendations until paired biological replicates for *P. copri* and *E. coli* were ready for mock community mixtures. At this point, final OD_600_ measurements for replicate cultures and previously calculated CFU/mL estimates were used to balance all cultures to an equal concentration of CFUs per mL by diluting in additional RNAlater.

Mock community mixtures were prepared on ice by mixing these equal concentration RNAlater suspensions as follows: (i) *P. copri* percentages of 100, 90, 50, 10, and 0 were prepared by mixing relative volumes of *P. copri* and *E. coli* suspensions to a total volume of 1 mL; (ii) 1% *P. copri* was prepared by mixing 100 µL of 1:10 diluted *P. copri* with 990 µL undiluted *E. coli;* and, (iii) 0.1% and 0.01% were prepared by mixing 100 and 10 µL of 1:100 diluted *P. copri* with 1000 µL undiluted *E. coli*. These mixtures were then pelleted by centrifuging (4 °C; 10,620 × *g*; 10 min). Pellets were frozen at −80 °C until nucleotide extraction was performed and sequencing libraries were prepared.

### Bacterial culture for *in vitro* cross-feeding experiments

Single colony monocultures of each isolate described in **Supplementary Fig. 8** were prepared from frozen stocks and acclimatized in PCDM and carbohydrate as described above. Acclimatization cultures (3 mL) were grown for 24 h and centrifuged (3,150 × *g*; 5 min) to pellet cells. Cells were then resuspended in PCDM without carbohydrate to remove any residual carbohydrate from acclimatization. All resuspensions had OD_600_ measured on a Tecan plate reader and were standardized to a blank-medium adjusted OD_600_ of 0.05. Five milliliter mono- and cocultures in anaerobically crimp-sealed culture tubes were prepared as follows: 5 mL of PCDM with 1% arabinan, arabinose, or glucose was added to each tube, then 50 μL of one organism for monocultures or 50 μL of each organism for cocultures were added. Cultures were sealed with rubber stoppers and crimp sealed, removed from the Coy chamber for baseline OD_600_ measurements and grown at 37 °C. OD_600_ was measured every 12 h, and triplicate cultures for each condition were collected every 24 h as follows: cultures were introduced into the Coy chamber, unsealed and transferred to 15 mL conical tubes, removed from the chamber, and centrifuged (3,150 × *g*; 5 min). For each resulting pellet, the supernatant was taken as 2 mL replicates of conditioned medium. The remaining medium was removed, and pellets were resuspended in 2 mL of RNAlater. These suspensions were split into 800 µL for DNA-RNA extraction and sequencing and 400 µL for phenol:chloroform genomic DNA extraction and qPCR, which were each centrifuged (10,620 × *g*; 10 min) before removal of RNAlater and storage at −80 °C.

### DNA and RNA sequencing

Ninety-six mock community bacterial samples described above were thawed from −80 °C, and DNA and RNA was extracted using the AllPrep DNA/RNA Kit (Qiagen; 80311). Barcoded DNA libraries were generated from raw DNA using the Nextera XT DNA Library Preparation Kit (Illumina; FC-131-1096). Barcoded Complementary DNA (cDNA) libraries were constructed from raw RNA using the Stranded Total RNA Prep with Ligation Kit with Ribo-Zero Plus Microbiome (Illumina; 20072063) for depletion of ribosomal RNA. DNA libraries were barcoded with IDT10 unique-dual indexes. cDNA libraries were barcoded with RNA UD Indexes Set D (Illumina; 20091661). Both DNA and cDNA libraries were sequenced initially using a MiniSeq instrument to balance pooled libraries to equal non-rRNA microbial-mapping reads as follows: raw reads were adapter-trimmed with a minimum read length of 50 bp using trimgalore (v0.6.4), filtered for reads that aligned to either the *Homo sapiens* GRCh38 or *Mus musculus* GRCm39 genome (bowtie2 v2.4.2), and used to quantify microbial gene abundances using kallisto (v0.46.2) using an index composed of predicted open reading frames from both the *P. copri* BgF5.2 and *E. coli* PS131 isolates^69,70^. The number of non-rRNA reads was calculated by totaling kallisto pseudocounts for genes and excluding counts from genes which were predicted as ribosomal RNAs by prokka (v1.14.6).

DNA libraries were sequenced on a NovaSeq X Plus instrument to a depth of 1.66 × 10^7^ ± 8.93 × 10^6^ paired-end 150 nt reads (mean ± s.d.). cDNA libraries were sequenced on NovaSeq X Plus instrument to a depth of 1.27 × 10^7^ ±2.61 × 10^6^ paired-end 150 nt reads. Sequencing reads were processed identically to above. After processing and quantification, pseudocounts assigned to ribosomal RNAs were removed to prevent uneven sequence removal by probe-based depletion from skewing library size normalization in downstream differential expression analyses. Pseudocounts were rounded to the nearest integer.

### Rarefaction of DNA and RNA counts

For DNA libraries, some samples were sequenced to a greater depth due to pool imbalance. For six samples that had a total raw depth greater than 2 × 10^7^ reads, we rarefied non-rRNA gene counts using the R package vegan implementation (vegan.rrarefy). Counts were randomly sampled without replacement to new sequencing depths randomly sampled from a normal distribution parameterized using the average and standard deviation number of microbial non-rRNA reads for the other 90 samples (12.3 ± 1.48 × 10^6^ reads). This resulted in 12.3 ± 1.46 × 10^6^ non-rRNA DNA reads per library across all 96 samples.

For cDNA libraries, given the imbalance of *P. copri* transcription rates and fractions of RNA reads between the arabinan and glucose conditions (**Supplementary Fig. 3b**), RNA reads from samples in the glucose condition were rarefied to balance the *P. copri* fraction of RNA reads between the conditions as follows. In cultures where glucose was the sole carbon source, *P. copri* reads were rarefied to depths randomly sampled from a normal distribution parameterized on the mean and standard deviation of the number of *P. copri* reads in cultures where arabinan was the carbon source. To maintain library size balance across samples and to account for under-sequencing of *E. coli* in the glucose samples due to *P. copri* overrepresentation in the pool of transcripts, additional *E. coli* counts for glucose samples were generated by rarefying from the six pure *E. coli* glucose samples (Pco0%). We rarefied *E. coli* reads to depths randomly sampled from a normal distribution parameterized using the mean and standard deviation number of *E. coli* reads in the corresponding arabinan samples. These additional *E. coli* counts were then combined with the existing counts from sequencing of each sample to generate profiles where both the total sequencing effort and the fraction of *P. copri* RNA reads were balanced between the arabinan and glucose conditions (**Supplementary Fig. 3d,e**). These rarefied RNA read profiles had 9.62 × 10^6^ ± 9.21 × 10^5^ non-rRNA microbial reads per library.

As described in **Supplementary Fig. 6**, benchmarking was carried out separately using the original RNA reads from each sample to test the ability of methods to report unbiased differential expression in the presence of confounding transcription rate changes between conditions. These original counts from cDNA libraries had 9.61 ± 1.12 × 10^6^ RNA reads per library.

### Validation analyses of DNA and RNA counts

For correlation analyses described comparing rarefied to original MTX count profiles (**Supplementary Fig. 3h-j**) and MTX counts produced with kallisto and bowtie2 (**Supplementary Fig. 3k-m**), sequencing effort was balanced across samples as follows. For species-level analysis for the rarefied and original MTX counts, scaling factors were calculated as the total non-rRNA RNA counts (depth) for a given organism in a given sample. Counts for all genes from each organism were linearly scaled such that each sample had the same depth for each organism. For comparisons of kallisto^74^ (alignment-free) and bowtie2^75^ (alignment-based) counts for each gene, correlations were performed after scaling counts in each sample to achieve equal total depth across organisms and to account for lower mapping rates of reads with bowtie2. Pearson correlation coefficients and simple ordinary least-squares linear regressions were then calculated between counts across all genes using scipy^89^. For probability distribution fitting (**Supplementary Fig. 3n,o**), DNA counts across all *P. copri* genes from all samples for each relative abundance and carbohydrate condition were used to fit gamma and log-normal distributions using scipy^91^. RNA counts were fit by estimating parameters for negative binomial and Poisson distributions from empirical mean and variance of counts data from all *P. copri* genes in the same samples.

### Differential expression method benchmarking using *in vitro* mock community datasets

#### Identification of differentially expressed genes

Metatranscriptomic counts from samples containing 100% *P. copri* grown in either arabinan or glucose were tested for differential expression using community-scaled DESeq2, with glucose treated as the reference condition. *P*-values and adjusted *P*-values of 0 were replaced with the minimum floating-point value for python (∼2×10^−308^). Statistical test results from DESeq2 were used to identify genes with differential expression under strict criteria (‘true positive’) by setting commonly used thresholds for |log_2_FoldChange| > 1 and adjusted *P*-value < 0.05. As noted in **Supplementary Table 4**, this resulted in 284 genes which were designated as upregulated in arabinan and 172 genes that were upregulated in glucose. Because many genes under this log_2_FoldChange threshold had significant *P*-values (**Fig. 3b**), we opted to use a stricter cutoff for defining ‘true negative’ genes to select those with strongest evidence for having no differential expression. A total of 854 ‘true-negative’ genes were identified by setting thresholds of |log2FoldChange| < 0.5 and adjusted P-value > 0.05. We elected to exclude genes that fell outside of these true positive and true negative sets (1731 genes) from performance metric calculations given their lack of strong evidence for or against differential expression when using DESeq2. In order to identify analogous gene sets using other differential expression methods (**Fig. 3c**), the same log fold-change and adjusted *P*-value criteria were applied to 100% *P. copri* test results produced by other packages. ‘Intersection’ was defined as the overlap between a method’s true positive and true negative gene sets and those defined by community-scaled DESeq2. ‘Disjoint’ was defined as the set of genes not present in the DESeq2 true positive/negative gene sets. A ‘consensus’ true positive set was defined as the union across all seven methods’ true positive genes identified from the monoculture samples; the same was done with true negative gene sets to obtain the consensus true negative set (see **Supplementary Fig. 5)**.

True positive rate was defined as the fraction of ‘true positive’ genes that had an adjusted P-value < 0.05 when tested by a given method. False positive rate was similarly defined as the fraction of ‘true negative’ genes that were erroneously tested as significantly different with an adjusted P-value < 0.05. Positive predictive value was defined as the fraction of positive test results (true and false positives with an adjusted *P*-value < 0.05) corresponding to true positive genes. ROC AUC was calculated using the sklearn implementation and imputing *P*-values of 1 for missing predictions while subsetting predictions to the explicitly defined true positive and true negative gene sets (i.e., excluding genes classified as ‘Other’ in **Fig. 3b**).

#### Benchmarking of existing methods

The same set of methods was applied as described in ‘*Differential expression method benchmarking using simulated datasets*.’ Methods were applied to each of the following sets of comparisons: those with no differential abundance, those with differential abundance and those with variable prevalence. For comparisons of equal abundance shown in **Fig. 3f,g**, the six samples for arabinan were compared against the six samples for glucose at each of the eight abundance levels. For comparisons with differential abundance shown in **Fig. 3h**, each pairwise comparison between the eight abundance levels in arabinan and in glucose were made, with the six corresponding samples for each carbohydrate condition and abundance being used as inputs. For comparisons with varying prevalence shown in **Fig. 3i**, for each of the eight abundance levels, the six samples for each carbohydrate condition had either 0, 2, 6 or 18 of the 0% *P. copri* (e.g., pure *E. coli*) samples added to each group to emulate prevalence levels of 100, 75, 50, and 25%. The *E. coli* samples added to each group were randomly sampled with replacement from the twelve 0% *P. copri* genes. For quantification of *E. coli* false positive rate shown in **Supplementary Fig. 6e**, this prevalence sampling scheme was modified to add in 0% *E. coli* (100% *P. copri*) samples. Non-colonized samples included non-zero DNA and RNA counts due to mis-mapping, a form of technical noise not seen in simulated datasets.

For 0% *P. copri* samples, imperfect mapping resulted in 5-28 DNA counts and 3-16 RNA counts assigned to *P. copri.* For 0% *E. coli* samples, imperfect mapping resulted in 3-713 DNA counts and 4-348 RNA counts assigned to *E. coli*. For all comparisons, glucose was defined as the reference condition, with positive log fold-change indicating upregulation in arabinan.

### Simulated analogs of mock communities

We applied MTX_synthetic, the previously published metagenomic and metatranscriptomic simulation framework^50^ that was used to create the above simulated datasets, to generate additional datasets which were more directly analogous to the mock communities analyzed in **Fig. 3**. We simulated two sets of additional datasets: (i) those with equal sample size (n=6 samples per condition) but equivalent in complexity to the original simulated datasets comprised of 100 species whose abundance and prevalence were based on Human Microbiome Project profiles, and (ii) those with equal sample size (n=6 samples per condition) comprised of two species whose relative abundances were based on the range of abundances for *P. copri* and *E. coli* in the mock communities. These new synthetic datasets are described and analyzed in **Supplementary Fig. 7**.

We made the following modifications to MTX_synthetic. First, we fixed an error where addition of a microbial feature whose abundance was equal to 1 minus the sum of the remaining microbial features caused invalid *P*-values to be supplied to numpy.random.multinomial. Second, we changed random assignment of samples to cases and controls to explicit splitting of the samples into cases (first half) and controls (second half) to ensure that each dataset had n=6 samples in each condition. Third, we seeded (i) random sampling of pangenomes for each organism, (ii) random selection of genes to be encoded for each genome and (iii) random selection of genes to be simulated as differentially expressed (‘spiked’ with expression) such that the same set of genes was simulated as differentially expressed (although with varying underlying DNA abundance and RNA expression levels) across all datasets to facilitate benchmarking comparisons. For more details on the simulation process, see *Supplementary Information* ref. 50.

For the first set of datasets with 100 species and 6 samples per group, the following parameters were used for all datasets: --n-bugs 100 --n-samples 12. For specific datasets, additional parameters were as follows: n12-true-exp-low: --spike-exp 0.1 --spike-exp-strength 0.50; n12-true-exp-med: --spike-exp 0.1 --spike-exp-strength 0.75; n12-true-exp: --spike-exp 0.1; n12-true-combo-dep-exp: --spike-dep --spike-exp 0.1; n12-true-combo-bug-exp: --spike-bug 0.5 --spike-exp 0.1.

For the second set of datasets mirroring mock community composition and abundances, the following parameters were used for all datasets: --n-bugs 2 --n-samples 12 --n-pangenes 3000 --n-groups 3000 --dna-read-depth 10500000 13500000 --rna-read-depth 8500000 10500000 --spike-exp 0.1. These were intended to mimic the number of species, samples, encoded and differentially expressed genes, and sequencing depths used for the mock communities.

Additionally, instead of the default HMP profile used to simulate empirical abundance and prevalence of human communities, custom HMP profiles were used that contained only *P. copri* and *E. coli* as microbial features. One such custom HMP profile was generated for each mixture level (e.g. Pco100%), with the corresponding relative abundances for *P. copri* populated using the 12 values of “Fraction of non-rRNA mapped reads assigned to *P. copri* genes” for rarefied MTX reads in **Supplementary Table 3b.** *E. coli* relative abundance was set as 1 – *P. copri* relative abundance.

The spiked set of genes for each set of datasets was used to calculate true positive rate, false positive rate, positive predictive value, negative predictive value, and ROC AUC as above (see ‘Differential expression method benchmarking using simulated datasets*’* above). Performance metrics were calculated using an alpha threshold of 0.05 for statistical significance of unadjusted *P*-values due to a low number of significant genes after multiple test correction for all benchmarked methods.

### Differential expression analysis of human metatranscriptomes

Methods used for generating 1,000 high quality metagenome-assembled genomes and filtering to 837 MAGs that met abundance and prevalence criteria are described in ref. 70. Quantification of metatranscriptomes was performed as described previously for 116 samples in the MDCF-2 treatment group at study weeks zero (pre-treatment) and 12 (end of treatment). Taxon-scaled DESeq2 was applied as implemented previously for analyses where all samples are included (**Fig. 5c-e**). An adjusted (across all genes, across all MAGs) *<u>P</u>*-value threshold of 0.25 was used for statistical significance.

#### Depth and detection filtering of samples

For a given genome with read counts for *m* genes in *n* samples, C_i,j_ describes the read count for gene *i* in sample *j*. For a given sample *j*, genome-level depth (*A_j_*) is defined as the sum of counts for all genes within a genome in that sample and gene-level detection is defined as the fraction of genes within a genome with detected (non-zero) counts in that sample. These are given in equations 1 and 2:

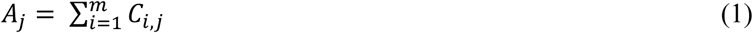

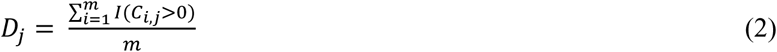

The inclusion of a sample under depth and detection filtering with a given depth threshold *α* and detection threshold *ß* is the product of indicator variables for its genome-level depth and gene-level detection exceeding the respective thresholds (equation 3):

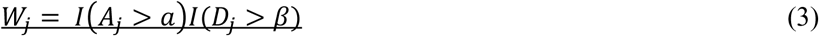

Samples were filtered individually for each MAG and were excluded if that MAG’s depth of sequencing coverage was less than the depth threshold (*α*) or if that MAG’s detection was less than the detection threshold (*ß*). MAGs were only advanced to differential expression testing if at least six samples in each treatment condition (timepoint) met filtering criteria and were not analyzed otherwise (**Supplementary Fig. 9b**).

For the parameter sweep described in **Supplementary Fig. 9**, we used depth thresholds of 0, 100, 1K, 10K, 100K, or 1M RNA reads and detection thresholds of 0%, 10%, 20%, 30%, 40%, 50%, 60%, 70%, 80%, or 90% to exclude samples for a given genome’s differential expression analysis. The number of differentially expressed genes was defined using a cutoff of 0.25 for adjusted *P*-values, and the number of MAGs with differential expression were similarly defined using the set of differentially expressed genes.

To identify genes whose differential expression was robust to sample exclusion, we considered the union of differentially expressed genes across each of the 60 sets of thresholds in the parameter sweep (yield 1,949 total genes). We opted to set strict criteria for defining genes robust to sample exclusion and required this set of genes to be present in at least 12 (20%) of the threshold pairs; we chose 12 threshold pairs to be greater than the number of possible values for both *α* and *ß,* such that genes in this set would have to be recovered at multiple values for both depth and detection thresholds. For comparisons of encoding genome-level depths in **Supplementary Fig. 9f**, genes in this set and the remainder of genes that were differentially expressed at least one time point had the depth (total RNA counts) of their encoding genomes averaged across all samples which were binned into the categories shown in the legend. For comparisons of the number of included samples in **Supplementary Fig. 9g**, the number of samples included for analysis for significantly differentially expressed genes across all threshold pairs were analyzed and split between the ≥12 threshold pairs set of genes and the remainder of differentially expressed genes.

### qPCR quantification of bacteria in cultures

Genomic DNA was extracted from mono- and coculture samples using a previously published phenol-chloroform extraction protocol^95^. Genomic DNA was diluted 1:100 in nuclease-free water to ensure concentrations within the linear range of the qPCR assay. qPCR assays were carried out using the QuantStudio 6 Flex system (Thermo Fisher Scientific) in 384-well format with 10 µL reactions. Each reaction contained 2.5 µL of 1:100 diluted genomic DNA, 0.36 µL each of 25 uM forward and reverse primers, 1.78 µL of nuclease-free water, and 5 µL of SYBR Select Master Mix (Fisher cat# 44-729-08). The cycling conditions were as follows: polymerase activation at 95 °C for 10 min; 40 cycles of 95 °C for 15 s, 56 °C for 15 s, 72 °C for 30 s; melting curve stage of 95 °C for 15 s, 60 °C for 60 s, 95 °C 15 s. The concentration of DNA from each organism was calculated using a 7-point 1:10 dilution series standard curve starting with 2 ng/µL genomic DNA obtained from monocultures of each organism in Wilkins-Chalgren broth. DNA concentrations were used to estimate genomic equivalents per mL in the original 5 mL cultures after accounting for dilutions and genome sizes (3.57 Mb for *P. copri* and 2.58 Mb for *M. multacida*), correcting for 16S rRNA copy number in each organism, and estimating the average molecular weight of a DNA base pair as 607.4 g/mol.

qPCR primers for *P. copri* targeting the 16S rRNA gene were used without modification from a previous publication^96^. qPCR primers for *Mitsuokella multacida* targeting the 16S rRNA gene were modified from a previously published assay targeting *Selenomonas ruminantium* and *Mitsuokella multacida*^97^ and checked for sequence specificity using BLAST. Primers were checked for specific amplification of the target bacteria and lack of activity on other organisms tested in coculture experiments. The primer sequences for each target were:

<u>Pco16S</u>

1. *P. copri* 16S rRNA F: 5’-CGCGAACTGGTTTCCTTGA-3’
2. *P. copri* 16S rRNA R: 5’-ACCGCTACACCACGAATTCC-3’
Mmu16S

1. *M. multacida* 16S rRNA 136-F: 5’-CTGCTAATACCGAATGTTGTGG-3’
2. *M. multacida* 16S rRNA 418-R: 5’-TACGAGTCGAAACCCTTCTTC-3’

### Mass spectrometry

Methods for extraction and quantification of amino acids were adapted from previous publications^70,98^. Briefly, aliquots of conditioned media were thawed, vortexed and 200 µL was mixed with 1 mL of methanol. Samples were vortexed, centrifuged (20,800 × *g*; 5 min) and 1 mL of the resulting supernatant was dried using a CentriVap (LabConco). The dried extract was resuspended in 100 µL of 10% methanol, centrifuged briefly, and a 80 µL aliquot was injected as input into an Agilent 1290 Infinity II ultra-high performance liquid chromatography system coupled with an Agilent 6470 Triple Quadrupole mass spectrometer operated in positive ion dynamic multiple reaction monitor mode.

## SUPPLEMETARY RESULTS

### Supplementary Note 1: Validation of mock community properties and methodology

DNA and RNA from these mock communities were extracted, sequenced, and mapped to the constituent genomes to produce gene-level counts for *P. copri* and *E. coli* (**Supplementary Table 3;** see *Methods*). The fraction of DNA reads mapping to *P. copri* genes was balanced between the arabinan and glucose growth conditions and approximated the emulated relative abundances (**Supplementary Fig. 3a**). The same was not true for the fraction of RNA reads; *P. copri* genes constituted a higher fraction of RNA reads in mixtures from glucose cultures compared to arabinan cultures (**Supplementary Fig. 3b**). This increase in *P. copri* counts was distributed genome-wide and not driven by a few genes; this suggests a globally increased transcription rate for *P. copri* in glucose, though we note that other explanations are also possible (e.g., decreased mRNA degradation, nucleotide extraction biases). Additionally, this increase was not detectable in 100% *P. copri* mixtures because sequencing counts are relative and not absolute abundances^52^. DESeq2 inferred far more differentially expressed genes upregulated in glucose in the 50% *P. copri* samples compared to the 100% *P. copri* samples (**Supplementary Fig. 3c**).

To characterize method performance, we initially opted to control for these global transcriptional changes by rarefying MTX counts from the *P. copri* in glucose samples to balance the fraction of *P. copri* reads between conditions (**Supplementary Fig. 3d**, see *Methods*). We elected to do this because recent analyses have demonstrated that rarefaction robustly controls for confounding differences in sequencing effort^71–73^. Furthermore, diversity analyses are much more sensitive to loss of low abundance features than differential expression (i.e., loss of features detected at low abundance can substantially reduce the estimated diversity of the whole sample). In differential expression analyses, highly expressed genes will retain sufficient quantification to infer differential expression whereas lowly expressed genes at the limit of detection likely would not have met criteria for statistical significance.

Similarly, we assumed most users aim to determine differential expression analogous to what is obtained from a single-organism sample, where inference of genes upregulated in each condition is not influenced by global transcription rate changes (**Fig. 1b,c**). Without rarefaction, the definition of ‘ground truth’ upregulated genes would otherwise change between the pure (100%) *P. copri* and mixed samples, which is both undesirable for benchmarking purposes and would lead to false positives when comparing against the reference single-organism comparison. While this confounding feature was removed from the datasets for initial benchmarking, transcription rate changes at the organism-level are a possibility in real-world datasets and may signify bacterial growth and death^39^ (see ‘*Benchmarking with confounding transcription rate changes*’ below).

To validate the rarefaction approach, we verified that total sequencing effort was balanced across samples and that differential expression was identified evenly in both carbohydrate conditions for rarefied but not original counts (**Supplementary Fig. 3e-g**). We then verified that rarefaction did not bias differential expression results by selectively depleting counts from individual genes. After correcting for changes in the total number of reads per organism, both *P. copri* and *E. coli* gene-level RNA counts were well correlated in a near 1:1 ratio between original and rarefied counts (**Supplementary Fig. 3h**). This generalized across samples where rarefaction was applied, indicating that gene-level quantifications relative to each organism’s total transcriptome were preserved (**Supplementary Fig. 3i,j**).

To verify that upstream processing of sequencing reads did not affect gene-level quantification, we also compared counts produced by an alignment-free tool, kallisto^74^, against an alignment-based tool, bowtie2^75^. After correcting for differences in total mapped counts due to bowtie2’s lower mapping rate, gene-level counts were well correlated, with a mean Pearson’s R of 0.993 and a mean linear regression slope of 1.03. (**Supplementary Fig. 3k-m**). Based on these analyses, we opted to use rarefied kallisto counts given more even fractions of *P. copri* transcripts between conditions and the increased number of pseudoaligned reads compared to bowtie2 counts. To determine suitable probability distribution models for DNA and RNA counts, we fit various distributions to the observed counts data (see *Methods*). The results revealed that non-zero DNA counts for all genes were well-approximated by the gamma distribution (**Supplementary Fig. 3n**). RNA counts were approximated by the negative binomial distribution, although probabilities for low but non-zero expressed genes were underestimated. (**Supplementary Fig. 3o**).

### Supplementary Note 2: Benchmarking with confounding transcription rate changes

We previously observed that *P. copri* genes constituted a higher fraction of RNA reads in mixtures from glucose cultures compared to those from arabinan cultures, resulting in biased inference of upregulation of genes in glucose (**Supplementary Fig. 3b,c**). Although this prevents a consistent definition of ‘ground truth’ gene upregulation between monocultures and mixtures, we leveraged this feature of the original sequencing counts from mock community datasets to determine whether statistical methods could correctly recover upregulated genes in both conditions despite confounding global transcription rate changes and produce results consistent with those obtained from sequencing a single-species sample. Given the larger fraction of RNA reads from *P. copri* in glucose, we additionally hypothesized that the compositional nature of sequencing data (i.e., the non-independence of features measured as relative quantities^80–81^) would lead to lower counts per gene for *E. coli*, resulting in false positives despite the lack of differences in *E. coli* growth conditions.

We repeated our previous benchmarking scheme and again used DESeq2 to define true positive and true negative gene sets in the 100% *P. copri* mixtures (**Supplementary Fig. 6a; Supplementary Table 6a,b**). We similarly defined 2,627 true negative genes from the 100% *E. coli* samples (**Supplementary Fig. 6b; Supplementary Table 6c**). The *P. copri* true positive and true negative gene sets were used to calculate a *P. copri*-specific true positive rate and false positive rate (P-TPR, P-FPR), while the *E. coli* true negative gene set was used to calculate an *E. coli*-specific false positive rate (E-FPR). The *P. copri* metrics quantify consistency of results with sequencing of a single-organism sample while the *E. coli* FPR measures false positives due to compositional shrinkage of *E. coli* expression measurements due to increased transcription by *P. copri* in glucose. As before, sensitivity for differentially expressed *P. copri* genes decreased at low relative abundances, though taxon-scaled methods better maintained sensitivity in this case (**Supplementary Fig. 6c; Supplementary Table 7a**). Taxon-scaled methods better controlled the *P. copri* FPR, indicating that they were able to recover results that were unbiased by the background changes in transcription rate (**Supplementary Fig. 6c**). Compositional effects resulted in an inflated *E. coli* FPR for all methods without taxon-scaling, though this was less apparent when *E. coli* comprised 99% or more of the sample and where compositional effects from *P. copri* transcription were modest due to its low abundance (**Supplementary Fig. 6c**).

In comparisons where *P. copri* had different relative abundance in the two conditions, sensitivity again decreased in the presence of differential abundance (**Supplementary Fig. 6d; Supplementary Table 7b**). Taxon-scaling controlled both *P. copri* and *E. coli* false positive rates in all differential abundance comparisons as long as the organism was not absent (0%) in either group, further indicating the ability of taxon scaling to adequately control for changes in transcription rate and compositional effects. Again, MTXmodel failed to infer true positive differential expression when confounding differential abundance was present, though it also exhibited fewer false positives for both organisms (**Supplementary Fig 6d**). In comparisons where *P. copri* had low prevalence (i.e., was not present in all samples), DESeq2 demonstrated decreased P-TPR, P-FPR, and E-FPR, while MPRAnalyze and MTXmodel were unaffected by inclusion of uncolonized samples (**Supplementary Fig. 6e; Supplementary Table 7c**).

### Supplementary Note 3: Validation of cross-feeding interactions *in vitro*

To validate this potential cross-feeding relationship identified in the mouse study, we advanced *P. copri* and *M. multacida* to *in vitro* mono- and coculture studies. Mono- and cocultures of *P. copri* and *M. multacida* were incubated in a defined medium^82^ supplemented with 1% (w/v) arabinan, arabinose, or glucose as the sole carbohydrate (n=3 replicate cultures/condition; see *Methods*). When grown on arabinan, *P. copri* monocultures exhibited modest growth achieving an OD_600_ ∼ 0.35 by 36 hours, while *M. multacida* alone did not demonstrate detectable growth. Coculture of the two exhibited increased total OD_600_ relative to each organism cultured independently (**Supplementary Fig. 8a; Supplementary Table 9**). In contrast, *P. copri* did not exhibit growth in glucose or arabinose in this time frame and cocultures of the two grew similarly to monocultures of *M. multacida*. We then used quantitative PCR (qPCR) to measure the abundance of each organism under each condition (n=3 biological replicates/condition; see *Methods*). Among substrates tested, only cocultures in arabinan supported simultaneous growth of both organisms and increased both *P. copri* and *M. multacida* cell density relative to either organism alone (**Supplementary Fig. 8b,c; Supplementary Table 10**). We advanced samples which exhibited detectable growth and yielded sufficient DNA and RNA to metagenomic and metatranscriptomic sequencing (**Supplementary Fig. 8d; Supplementary Table 11, 12**).

We then performed differential expression analysis to determine whether transcriptional changes by *M. multacida* in response to arabinan cross-feeding recapitulated the *in vivo* response to *P. copri* colonization. In simple comparisons of pure *Mitsuokella* samples grown in arabinose vs. glucose, MTXmodel and both the community-scaled and taxon-scaled DESeq2 approaches successfully inferred the expected upregulation of arabinose utilization genes in the pentose phosphate pathway (**Supplementary Fig. 8e; Supplementary Table 13**). The same differential expression of genes involved in glutamate, tryptophan, and glutamine biosynthesis observed *in vivo* were also detected by both DESeq2 approaches but not with MTXmodel. In comparisons of arabinan cocultures with glucose controls and along the time course of the experiment, MTXmodel provided high variance fold-change estimates and did not infer any significant genes in these pathways or the entire *M. multacida* genome (**Supplementary Fig. 8f,g**). DESeq2 detected expected upregulation of arabinose utilization genes in both comparisons and additionally inferred upregulation of tryptophan, glutamate, and glutamine biosynthesis at 72 h compared to 24 h (**Supplementary Fig. 8f,g**). Taxon-scaling resulted in relatively even differential expression inference for both conditions in the arabinan versus glucose comparison while community-scaling inferred a distribution of log-fold changes skewed towards upregulation in glucose, consistent with the differential abundance of *M. multacida* between the mono- and cocultures (violin plots, **Supplementary Fig. 8f**).

We applied targeted mass spectrometry to conditioned media samples to quantify tryptophan, glutamate, and glutamine and to determine whether transcriptional changes were associated with their availability. Compared to the *P. copri* monocultures in arabinan, the cocultures produced significantly more glutamate and depleted significantly more glutamine and tryptophan at 72 hours, indicating that *M. multacida* upregulation of these biosynthesis pathways was associated with low availability of tryptophan and glutamine and increased availability of glutamate (**Supplementary Fig. 8h**). In contrast, amino acid levels were not significantly different between endpoint cocultures and *M. multacida* monocultures for either monosaccharide condition (**Supplementary Fig. 8h**). Based on these results, we concluded that taxon-scaled DESeq2 was able to nominate cross-feeding interactions and downstream amino acid metabolic responses from our gnotobiotic animal studies which could then be validated *in vitro*.

---

**Supplementary Figure 1.**
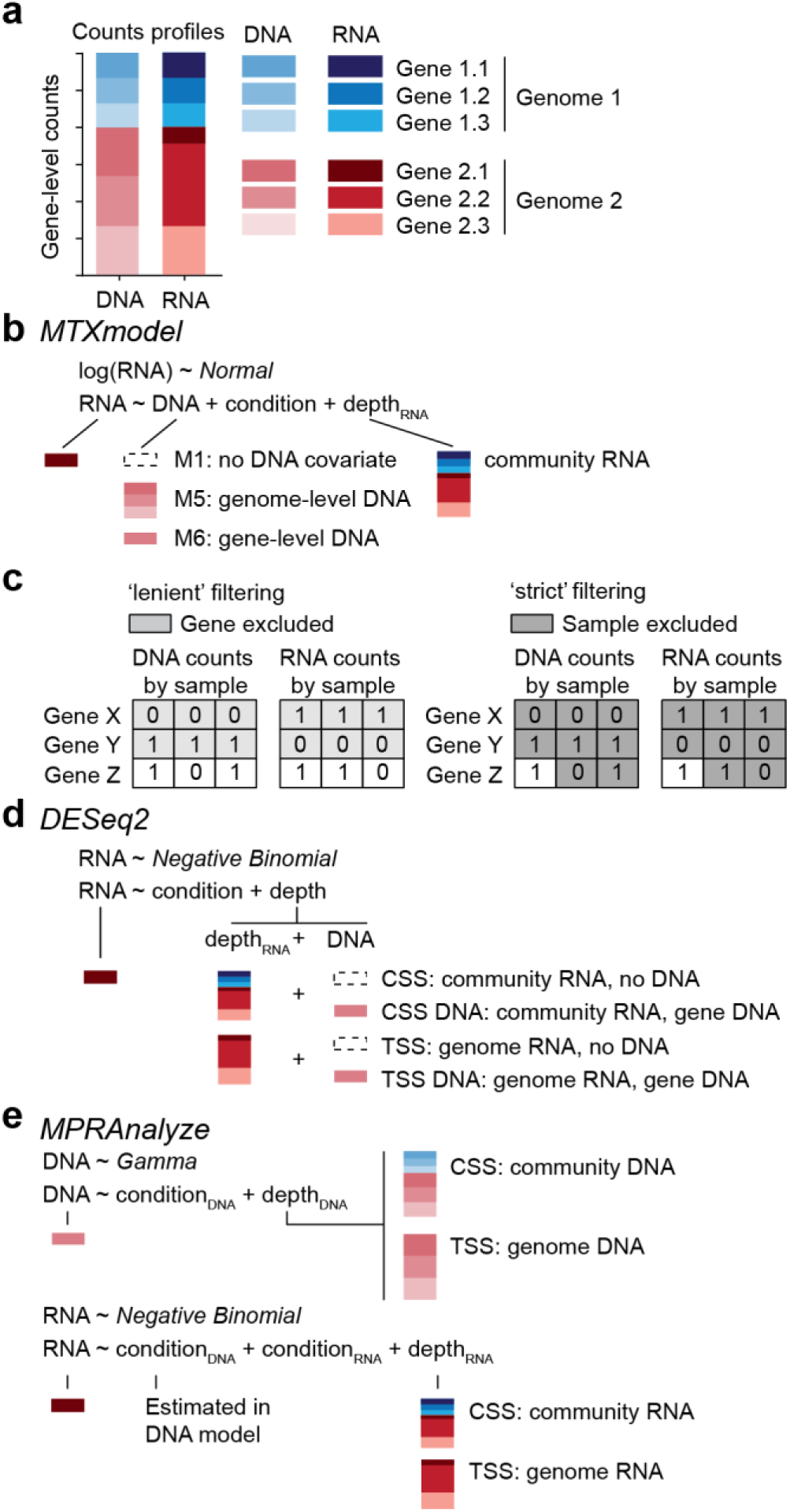
Metatranscriptomic differential expression statistical models. **(a)** Profiles of DNA and RNA counts for a sample containing two genomes (red and blue) with three genes each. **(b)** Statistical model used by MTXmodel under models M1, M5, and M6. **(c)** Zero-filtering approaches used by MTXmodel with ‘lenient’ gene-level filtering or ‘strict’ sample-level filtering. **(d)** Statistical model used by DESeq2 with either community- (CSS) or taxon-scaling (TSS) as well as with or without DNA covariates (DNA). **(e)** Statistical model used by MPRAnalyze for both DNA and RNA counts under community- and taxon-scaling. Abbreviations: M1, no DNA covariate; M5, genome-level DNA covariate; M6, gene-level DNA covariate; CSS, community-sum-scaling; TSS, taxon-specific-scaling. CSS DNA, community-scaling with DNA abundance normalization; TSS DNA, taxon-scaled with DNA abundance normalization.

**Supplementary Figure 2.**
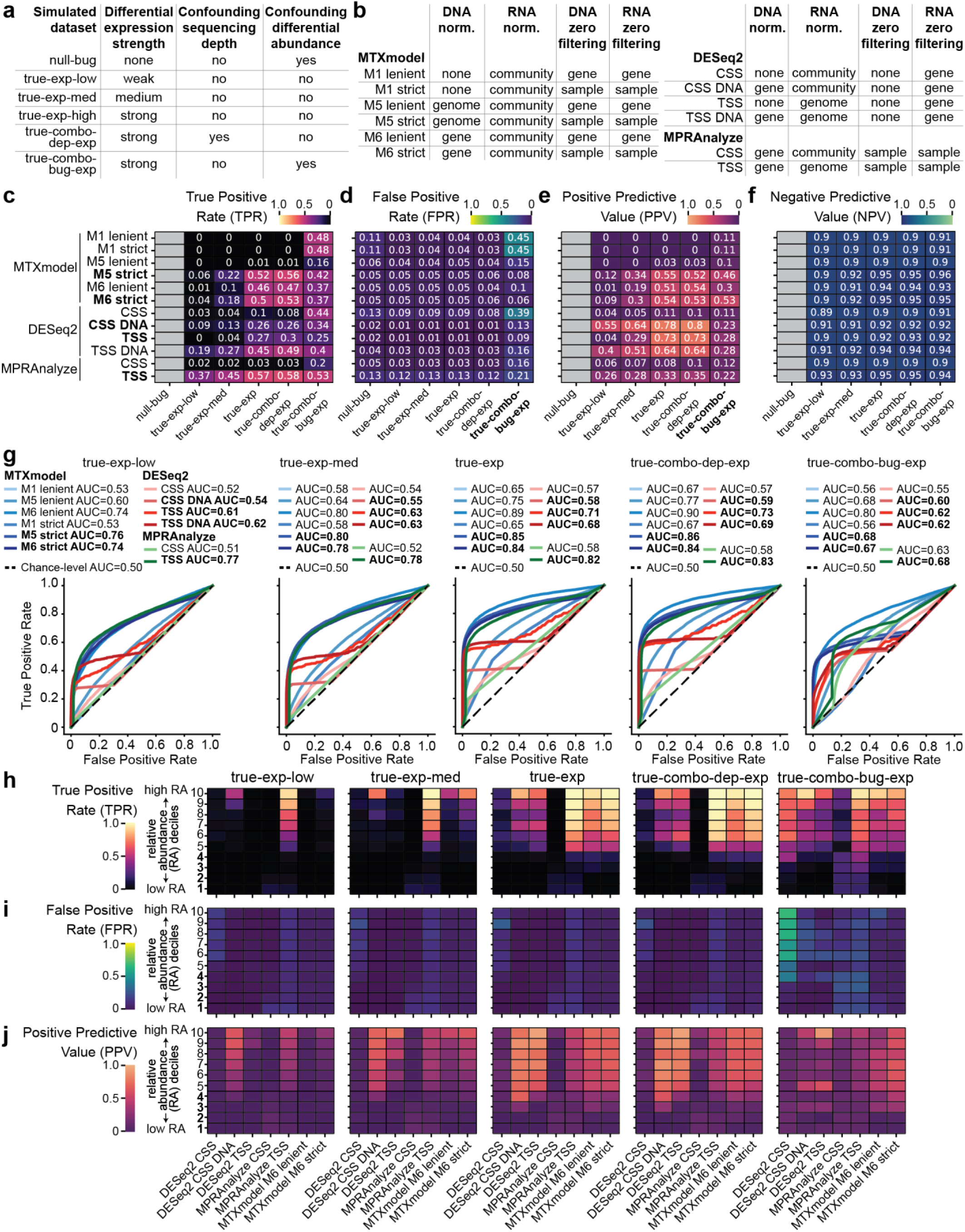
Statistical differential expression method benchmarking on simulated datasets for all methods tested. **(a,b)** Characteristics of simulated datasets (panel a) and differential expression methods and implementations tested (panel b). **(c-f)** True positive rate (TPR, or sensitivity; panel c), false positive rate (FPR, or 1-specificity; panel d), positive predictive value (PPV, or precision; panel e), and negative predictive value (NPV; panel f) for MTXmodel, DESeq2, and MPRAnalyze across a range of parameter choices in six simulated datasets. Implementations and datasets specifically emphasized in the text are bolded. Metrics that are not defined for datasets lacking ground truth differential expression are grayed out. (**g**) Area under the curve (AUC) quantification and receiver operating characteristic curves for statistical methods in the five synthetic ‘true-’ datasets that included simulated differential expression. An AUC of 0.5 indicates the classification power of random guessing, with greater AUC indicating a greater ability to discriminate differentially expressed and non-differentially expressed genes. (**h-j**) True positive rate (panel h), false positive rate (panel i), and positive predictive value (panel j) for a subset of methods when applied to gene sets originating from organisms ranked and binned by their relative abundance (RA) in the simulated datasets. Abbreviations: M1, no DNA abundance covariates; M5, genome-level DNA covariates; M6, gene-level DNA covariates; CSS, community-sum-scaling; TSS, taxon-specific-scaling; DNA, DNA abundance normalization.

**Supplementary Figure 3.**
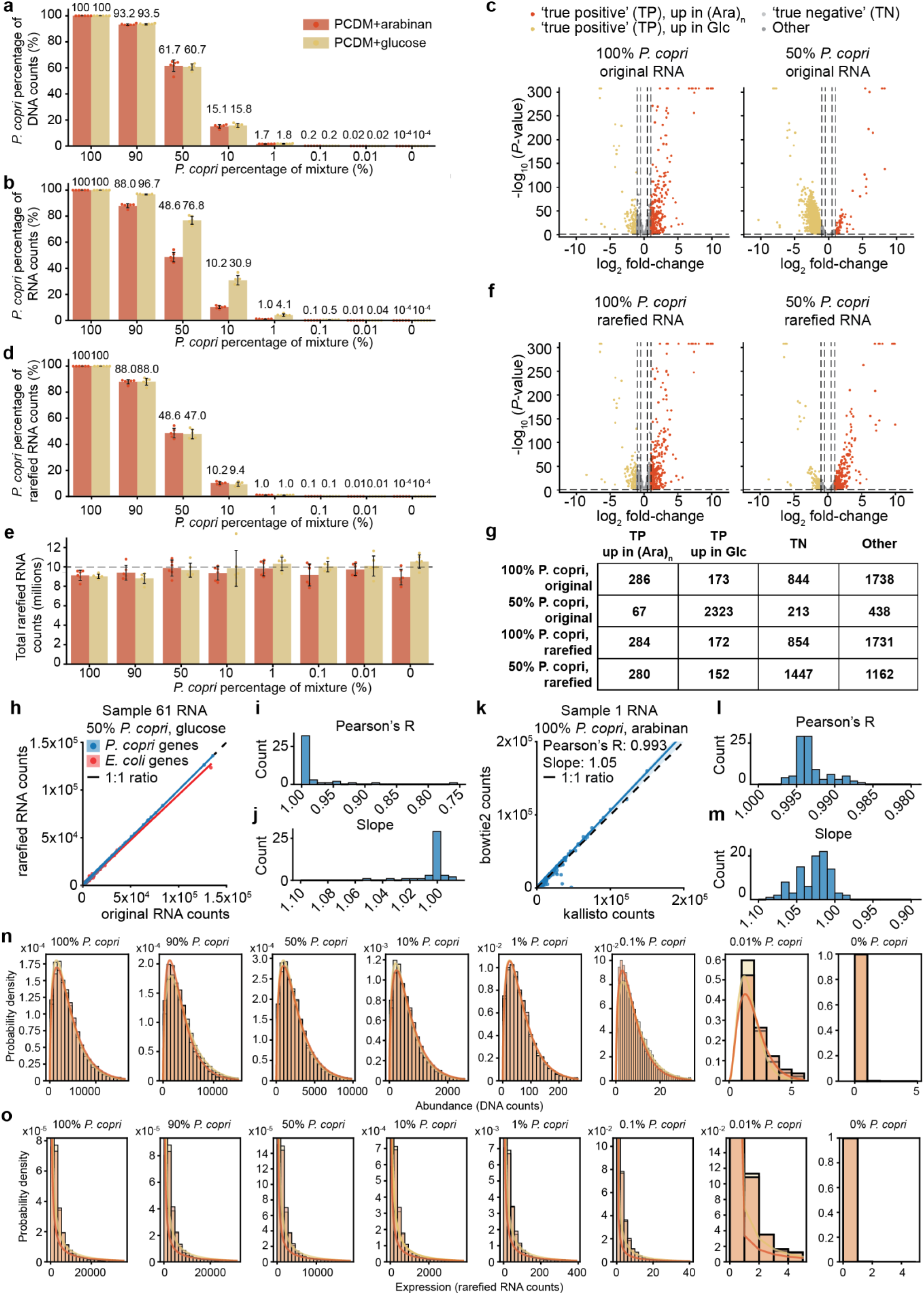
Validation and characterization of mock communities. **(a,b)** Percentage of non-rRNA microbial DNA (panel a) and RNA (panel b) reads mapping to *P. copri* for each mixture ratio and each condition. Mean ± standard deviation values are shown for the six replicates in each condition. **(c)** Volcano plots for results generated by community-scaled DESeq2 when applied to the 100% vs. 50% *P. copri* samples, demonstrating how increases in total transcription rate skew differential expression testing but are not readily detected in single-organism samples. (**d,e**) Percentage of non-rRNA microbial RNA reads mapping to *P. copri* (panel d) and total microbial non-rRNA reads (panel e) in mock community counts profiles after applying rarefaction to balance *P. copri* sequencing effort between conditions. (**f**) Volcano plots generated by community-scaled DESeq2 from rarefied counts profiles, indicating correction of varying skewness of differential expression results across abundance levels. (**g**) Number of genes meeting true positive (TP) and true negative (TN) log fold-change and adjusted *P*-value thresholds for 100% and 50% *P. copri* comparisons using the original and rarefied counts. Directionality and number of TP genes are maintained between datasets using rarefied but not original counts. (**h**) Representative regression plot of *P. copri* (blue) and *E. coli* (red) gene-level RNA counts after adjusting for total counts for each organism for one sample containing 50% of each organism. The dashed black line shows a 1:1 ratio, indicating preservation of gene-level relative representation within each organism’s total transcriptome. (**i,j**) Histograms of Pearson’s correlation coefficients (Pearson’s R; panel i) and ordinary least squares linear regression slopes (panel j) comparing gene-level RNA counts after adjusting for organism-level sequencing effort (as in panel h) across the 48 glucose-conditioned samples where rarefaction was applied. **(k)** Representative regression plot of kallisto pseudoaligned counts vs. bowtie2 aligned counts for all *P. copri* genes in one sample after adjusting for total mapped reads to account for the lower mapping rate of bowtie2. Pearson’s correlation coefficient and the slope of a simple linear model are annotated. **(l,m)** Histograms of Pearson’s correlation coefficients (panel l) and linear model slopes (panel m) for correlations between kallisto and bowtie2 counts for all 96 samples. (**n,o**) Non-zero DNA counts (panel n) and all RNA counts (panel o) across genes for each of the six replicate samples for each mixture ratio and carbohydrate condition. The probability density function for gamma distributions fit by maximum likelihood estimation for each condition are shown as solid lines in panel n. The probability mass functions of negative binomial distributions fit by the method of moments are shown as solid lines in panel o. PCDM, *P. copri* defined medium, adapted from a previous publication^82^.

**Supplementary Figure 4.**
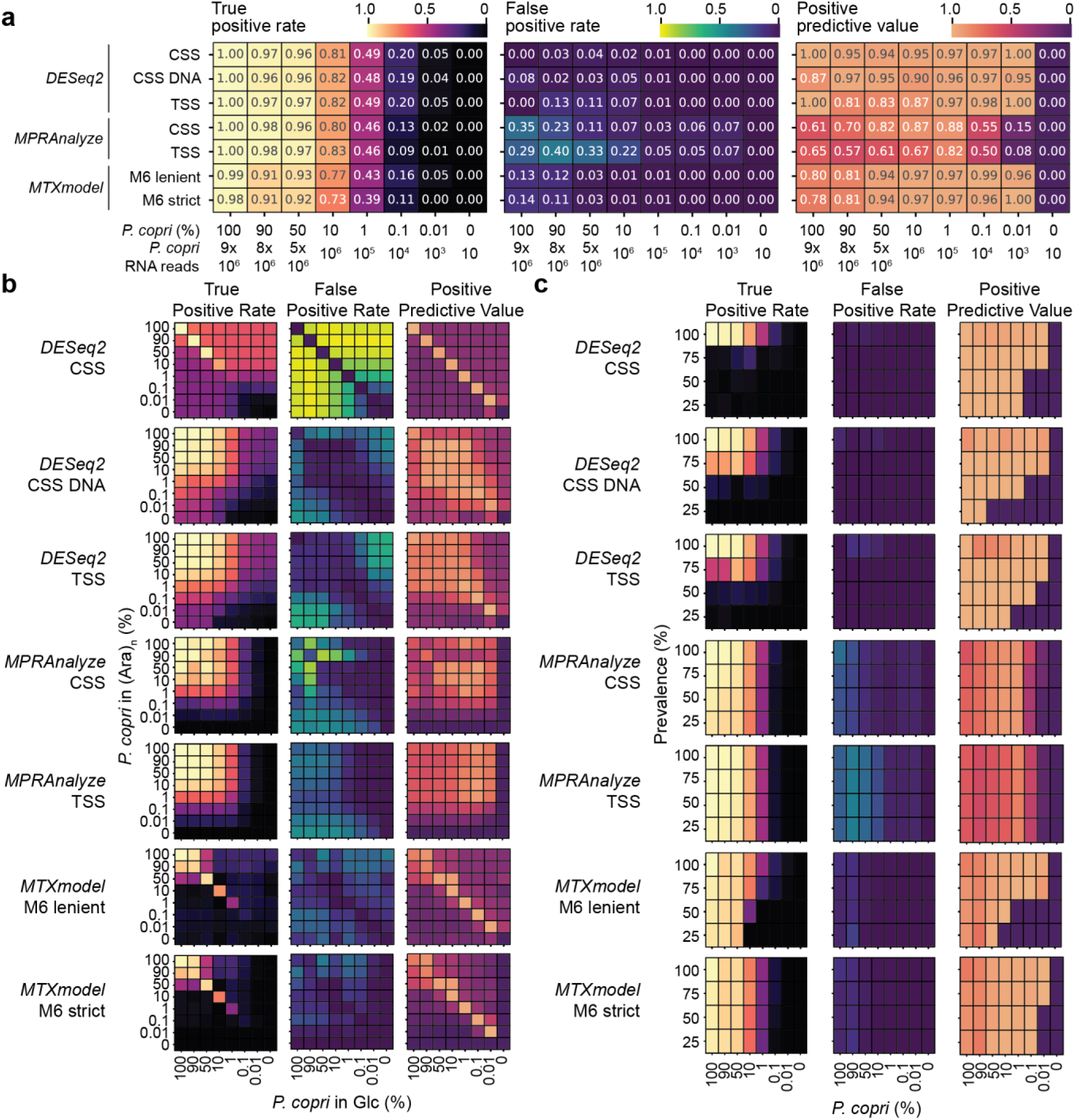
Sensitivity, specificity, and precision of datasets from mock communities. (**a**) True positive rate (TPR), false positive rate (FPR), and positive predictive value (PPV) for benchmarked methods when analyzing the different mixture ratios of *P. copri*, from highest to lowest relative abundance. (**b**) TPR, FPR, and PPV for methods in comparisons with varying strengths of differential abundance, emulated by performing all pairwise comparisons between mixture ratios. (**c**) TPR, FPR, and PPV of methods when emulating decreasing prevalence by including either 0, 2, 6, or 18 100% *E. coli* samples with the six *P. copri*-containing samples in each group. Abbreviations: (Ara)_n_, arabinan; Glc, glucose; TP, true positive; TN, true negative; CSS: community-sum-scaling; CSS DNA: community-sum scaling with DNA abundance normalization; TSS, taxon-specific-scaling; TPR, true positive rate; FPR, false positive rate; PPV, positive predictive value.

**Supplementary Figure 5.**
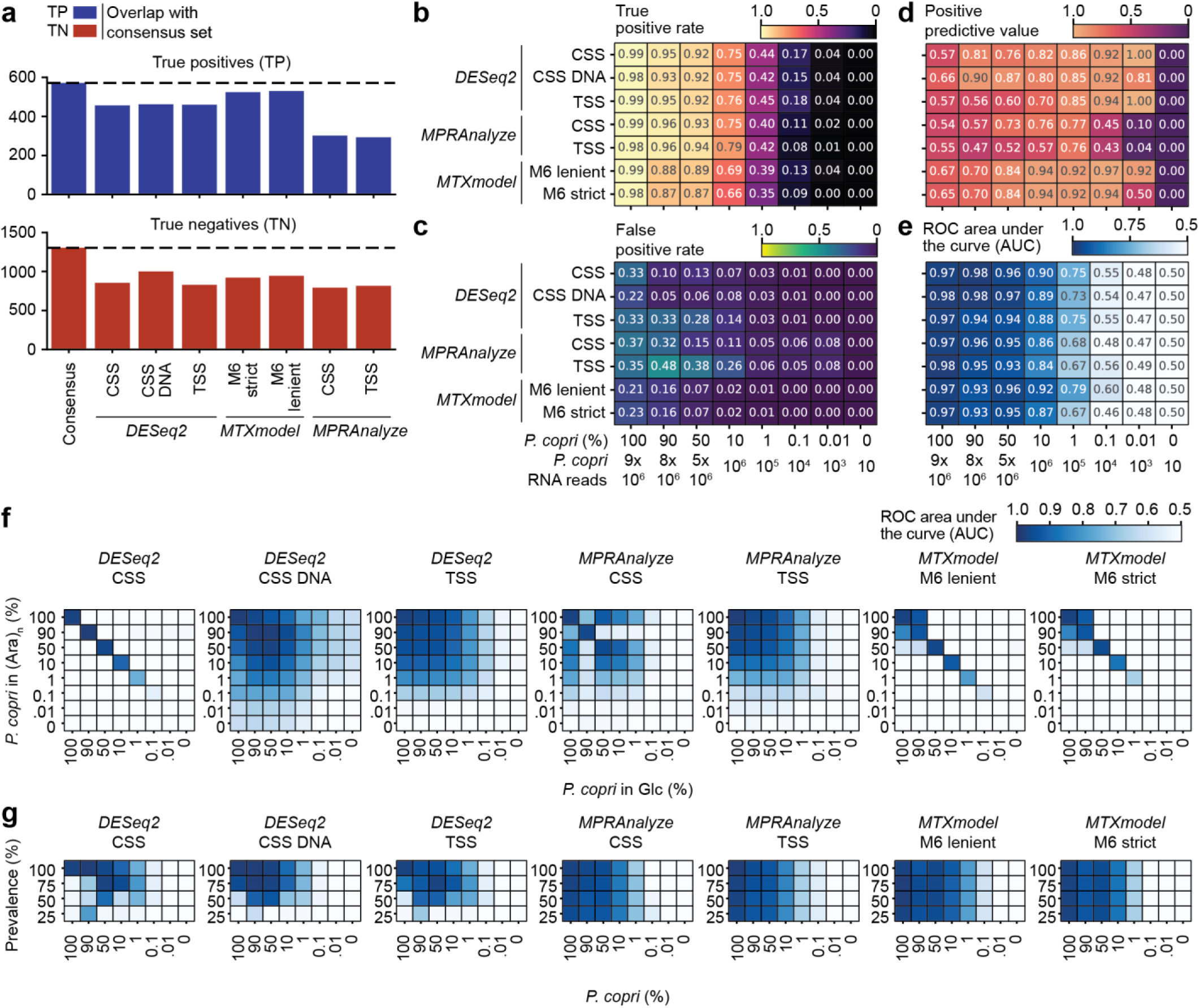
Benchmarking on mock communities using consensus sets of true positive and true negative genes. **(a)** Consensus true positive genes (n=571) and true negative genes (n=1,304) were defined as the union across the true positive and true negative gene sets determined using individual methods in Fig. 3c. Dashed lines indicate the size of the consensus sets and each method’s intersection with the consensus sets are shown in colored bars. (**b-e**) True positive rate (panel b), false positive rate (panel c), positive predictive value (panel d), and ROC AUC (panel e) calculated using the consensus gene sets for benchmarked methods when analyzing the different mixture ratios of *P. copri*, from highest to lowest relative abundance. (**f,g**) ROC AUC quantification in comparisons confounded by differential abundance (panel f) and varying prevalence of *P. copri* (panel g). Abbreviations: TP, true positive; TN, true negative; CSS: community-sum-scaling; CSS DNA: community-sum scaling with DNA abundance normalization; TSS, taxon-specific-scaling; TPR, true positive rate; FPR, false positive rate; PPV, positive predictive value; ROC, receiver operating characteristic; AUC, area under the curve quantification.

**Supplementary Figure 6.**
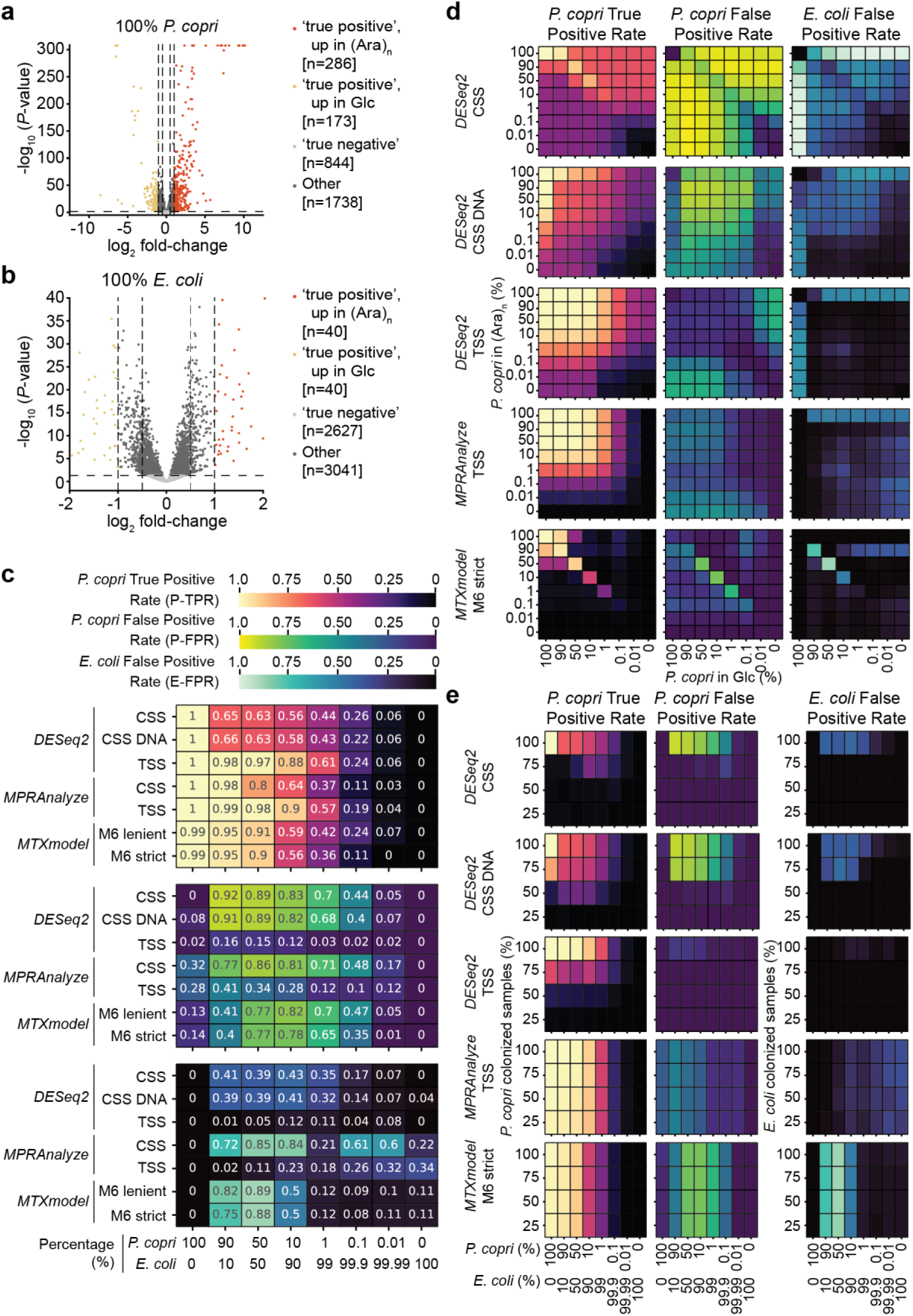
Benchmarking on datasets from mock communities with confounding transcription rate changes and compositional effects. **(a,b)** True positive and true negative gene sets defined by DESeq2 for 100% *P. copri* samples (panel a) and for 100% *E. coli* samples (panel b). (**c**) True positive rate for the *P. copri* TP gene set (P-TPR), false positive rate for the *P. copri* TN gene set (P-FPR), and false positive rate for the *E. coli* TN gene set (E-FPR) for each of the methods for each mixture ratio, in order of descending *P. copri* relative abundance. (**d**) P-TPR, P-FPR, and E-FPR for datasets with emulated differential abundance for DESeq2 with community- or taxon-scaling, MPRAnalyze with taxon-scaling, and MTXmodel using gene-level DNA covariates. (**e**) P-TPR and P-FPR for datasets with emulated non-colonization by inclusion of 100% *E. coli* samples in each group. Abbreviations: CSS, community-sum-scaling; CSS DNA, community-sum scaling with DNA abundance normalization; TSS, taxon-specific-scaling; P-TPR, *P. copri* true positive rate; P-FPR, *P. copri* false positive rate; E-FPR, *E. coli* false positive rate.

**Supplementary Figure 7.**
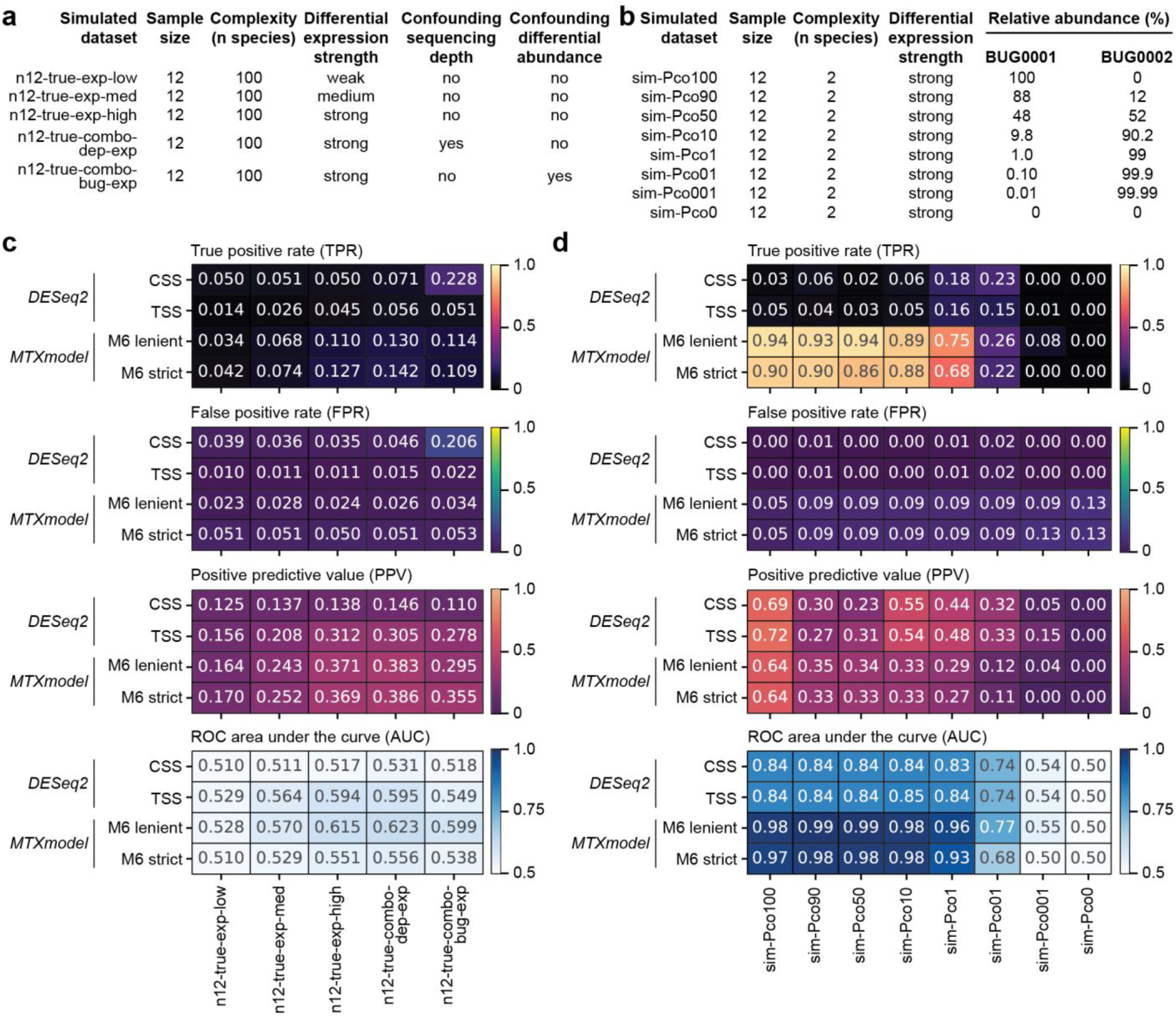
Differential expression method benchmarking on simulated datasets with analogous properties to mock communities. **(a,b)** Characteristics of simulated datasets with sample size identical to mock community datasets (panel a) and simulated datasets with both sample size, complexity, and relative abundances based on mock communities (panel b). (**c,d**) True positive rate, false positive rate, and positive predictive value calculated at a significance threshold of 0.05, as well as ROC AUC calculated across significance thresholds for the 100 species synthetic communities (panel c) and the two-member synthetic communities (panel d). Abbreviations: CSS: community-sum-scaling; TSS, taxon-specific-scaling; TPR, true positive rate; FPR, false positive rate; PPV, positive predictive value; ROC, receiver operating characteristic; AUC, area under the curve.

**Supplementary Figure 8.**
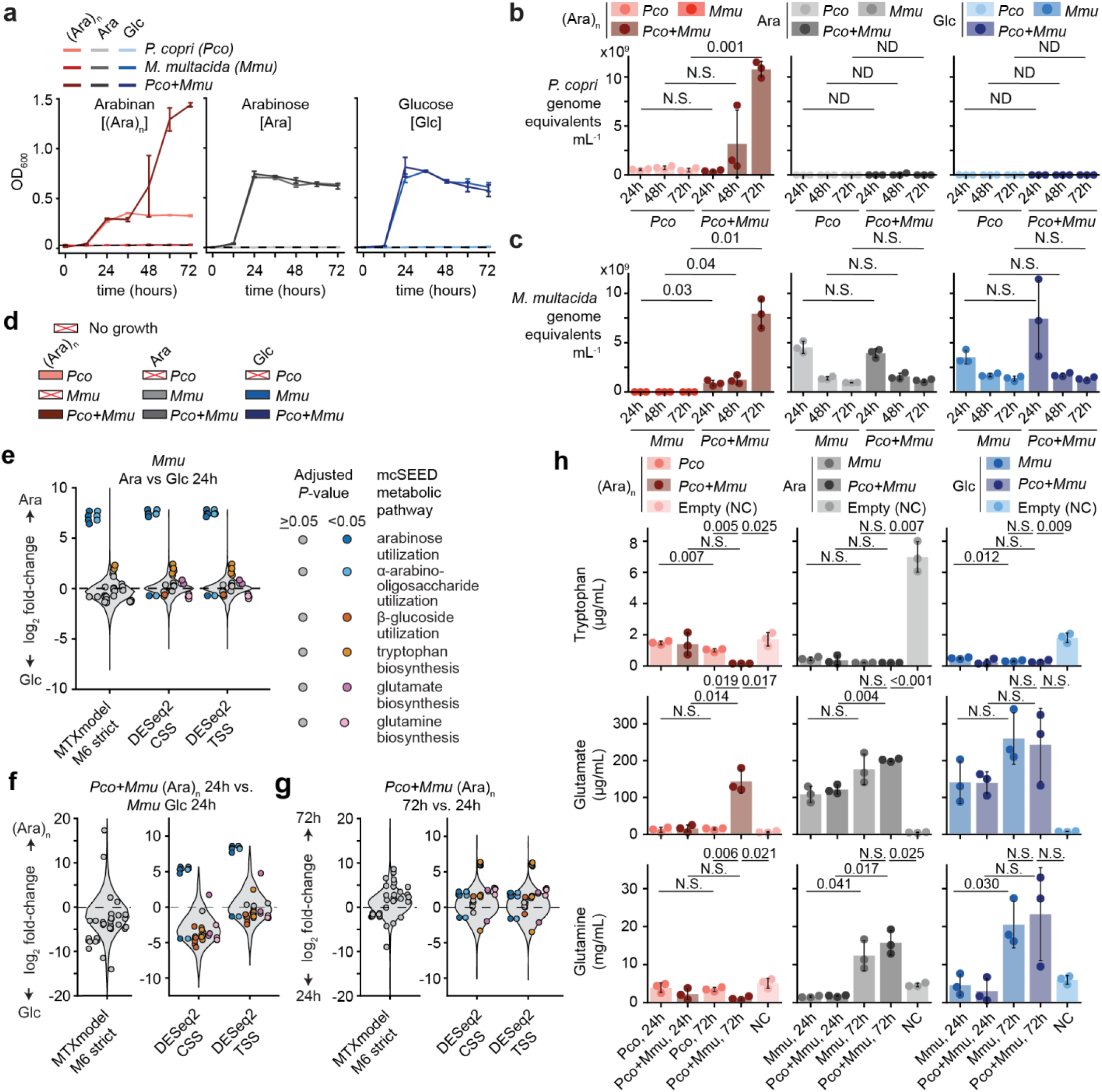
*In vitro* validation of arabinan cross-feeding between *P. copri* and *M. multacida*. **(a)** Growth of monocultures and cocultures of *<u>P. copri</u>*(*Pco*) and *M. multacida* (*Mmu*) in PCDM supplemented with either 1% arabinan, arabinose, or glucose. **(b,c)** qPCR quantification of *P. copri* (panel b) and *M. multacida* (panel c) in mono-and cocultures. **(d)** Experimental design for samples advanced to sequencing and mass-spectrometry based on growth and qPCR quantification in panels a-c. **(e-g)** Differential expression results for *M. multacida* genes predicted to be involved in arabinose, alpha-arabinoside, and beta-glucoside utilization, as well as biosynthesis of glutamate, glutamine, and tryptophan as reported by the three methods using an adjusted *P*-value threshold of 0.05. Violin plots represent the distribution of log2 fold-change across all genes in the *M. multacida* genome; points represent differential expression test results for individual genes from select metabolic pathways, horizontally grouped by pathway and colored if they were significantly differentially expressed with a given method. The dashed line indicates a fold-change of 1 (no change in expression). Comparisons between arabinose and glucose *M. multacida* monocultures are shown in panel e, comparisons between arabinan cocultures and *M. multacida* in glucose are shown in panel f, and comparisons between 72 h and 24 h arabinan cocultures are shown in panel g. **(h)** Mass-spectrometric quantification of glutamate, glutamine, and tryptophan from conditioned media collected from mono- and cocultures every 24h for each condition. Mean ± standard deviation values are shown in panels a-c and h. *P*-values were calculated by a two-sided Welch T-test of independent samples with unequal variance in panels b, c, and i, and the specified differential expression methods in panels e-g. Abbreviations: (Ara)n, arabinan; Ara, arabinose; Glc, glucose; *Pco*, *P. copri*; *Mmu*, *M. multacida*; N.S., not statistically significant (Benjamini-Hochberg adjusted P-value > 0.1); ND no detection in both groups (quantification lower than inoculum input); NC, no inoculum negative control.

**Supplementary Figure 9.**
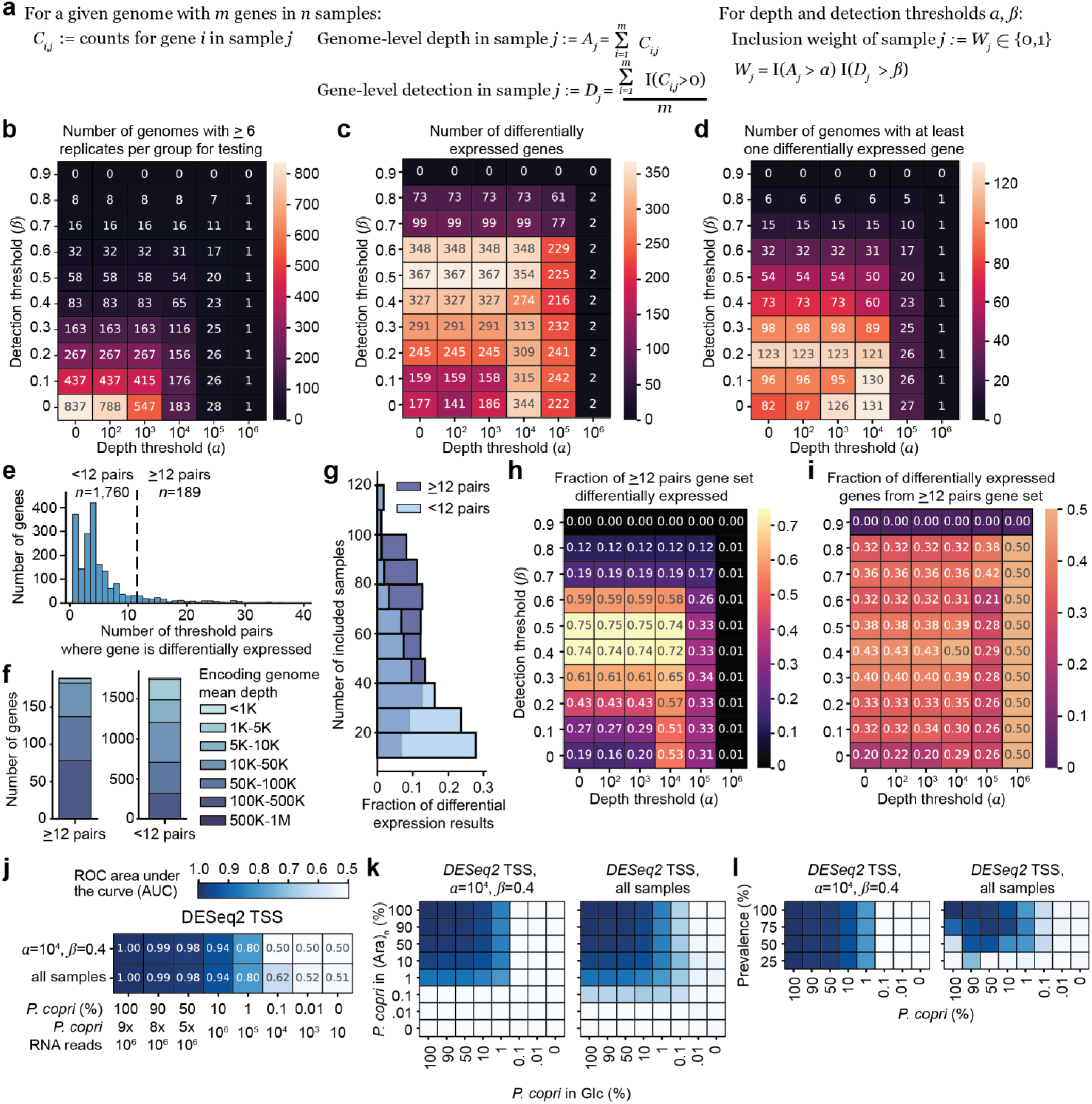
Parameter sweep of genome-level depth and gene-level detection thresholds for sample inclusion in differential expression analysis of the human study. **(a)** Definition of genome-level depth and gene-level detection metrics for a given sample, as well as inclusion of a sample at depth and detection thresholds *α* and *ß*. Samples are filtered separately for each genome’s differential expression analysis if their genome-level depth is not greater than *α* or if their gene-level detection is not greater *ß*. **(b-d)** The number of genomes with sufficient samples for testing for differential expression (panel b), the number of differentially expressed genes (panel c), and the number of genomes with at least one differentially expressed gene (panel d) across a range of depth (*α*) and detection thresholds (*ß*). **(e)** The distribution of the number of threshold pairs where a gene was significantly differentially expressed for all genes which were significant in at least one set of thresholds (1,949 genes). A set of 189 genes was defined as having inferred differential expression in at least 12 (20%) of the threshold pairs. **(f,g)** The average depth of genomes encoding genes (panel f) and the number of samples included for differential expression analysis (panel g) for genes recovered in at least 12 threshold pairs or fewer than 12 threshold pairs. **(h,i)** The fraction of this set of genes which was inferred as differentially expressed (panel h) and the fraction of all significant differential expression results from this set of genes (panel i) across the range of depth (*α*) and detection thresholds (*ß*). Statistical significance was defined using a cutoff of 0.25 for *P*-values adjusted across all genes from all genomes with sufficient replicates for analysis (up to 1,929,056 genes with no sample filtering). (**j-l**) ROC AUC for differentially expressed genes in mock communities in comparisons with no differential abundance (panel j), differential abundance (panel k), or low prevalence (panel l) for taxon-scaled DESeq2 either including all samples or filtering samples using depth and detection thresholds of *α*=10^4^ *and ß*=0.4. Abbreviations: ROC, receiver operating characteristic; AUC, area under the curve; TSS, taxon-specific-scaling.

